# Systems genetic dissection of Alzheimer’s disease brain gene expression networks

**DOI:** 10.1101/2024.10.04.616661

**Authors:** Pinghan Zhao, Omar El Fadel, Anh Le, Carl Grant Mangleburg, Justin Dhindsa, Timothy Wu, Jinghan Zhao, Meichen Huang, Bismark Amoh, Aditi Sai Marella, Yarong Li, Nicholas T. Seyfried, Allan I. Levey, Zhandong Liu, Ismael Al-Ramahi, Juan Botas, Joshua M Shulman

**Affiliations:** Department of Neuroscience, Baylor College of Medicine, Houston, TX 77030, USA; Jan and Dan Duncan Neurological Research Institute, Texas Children’s Hospital, Houston, TX, USA; Department of Molecular and Human Genetics, Baylor College of Medicine, Houston, TX 77030, USA; Medical Scientist Training Program, Baylor College of Medicine, Houston, TX, 77030, USA; Department of Neurology, Baylor College of Medicine, Houston, TX 77030, USA; Department of Biochemisry, Emory University, Atlanta, GA 30329; Department of Neurology, Emory University, Atlanta, GA 30329; Center for Neurodegenerative Disease, Emory University, Atlanta, GA 30329; Department of Pediatrics, Baylor College of Medicine, Houston, TX 77030, USA; Center for Alzheimer’s and Neurodegenerative Diseases, Baylor College of Medicine, Houston, TX, 77030, USA

**Keywords:** neurodegenerative disease, transcriptome, coexpression, excitotoxicity, systems biology, glutamate, innate immune, neuroinflammation

## Abstract

In Alzheimer’s disease (AD), changes in the brain transcriptome are hypothesized to mediate the impact of neuropathology on cognition. Gene expression profiling from postmortem brain tissue is a promising approach to identify causal pathways; however, there are challenges to definitively resolve the upstream pathologic triggers along with the downstream consequences for AD clinical manifestations. We have functionally dissected 30 AD-associated gene coexpression modules using a cross-species strategy in fruit fly (*Drosophila melanogaster*) models. Integrating longitudinal RNA-sequencing and behavioral phenotyping, we interrogated the unique and shared transcriptional responses to amyloid beta (Aβ) plaques, tau neurofibrillary tangles, and/or aging, along with potential links to progressive neuronal dysfunction. Our results highlight hundreds of conserved, differentially expressed genes mapping to human AD regulatory networks. To confirm causal modules and pinpoint AD network drivers, we performed systematic *in vivo* genetic manipulations of 357 conserved, prioritized targets, identifying 141 modifiers of Aβ- and/or tau-induced neurodegeneration. We discover an up-regulated network that is significantly enriched for both AD risk variants and markers of immunity / inflammation, and which promotes Aβ and tau-mediated neurodegeneration based on fly genetic manipulations in neurons. By contrast, a synaptic regulatory network is strongly downregulated in human brains with AD and is enriched for loss-of-function suppressors of Aβ/tau in *Drosophila*. Additional experiments suggest that this human brain transcriptional module may respond to and modulate Aβ-induced glutamatergic hyperactivation injury. In sum, our cross-species, systems genetic approach establishes a putative causal chain linking AD pathology, large-scale gene expression perturbations, and ultimately, neurodegeneration.

## INTRODUCTION

Alzheimer’s disease (AD) is rapidly increasing in prevalence due to population aging and remains neither curable nor preventable^1^. A major goal for the field is to map the AD pathophysiologic cascade from its neuropathologic triggers to the downstream cognitive manifestations (Fig. 1a). Gene expression profiling from human brains is one promising approach to highlight molecular mediators. In a meta-analysis of RNA-Sequencing (RNAseq) from ∼2,000 postmortem brain tissue samples, the Accelerating Medicines Partnership (AMP)-AD target discovery consortium defined 30 AD-associated consensus coexpression networks^2–5^. This study highlighted robust transcriptional signatures for immune and synaptic regulatory mechanisms in AD pathogenesis, among other pathways, and similar findings have been independently reported by others^6–11^. Additional evidence from human genome-wide association studies implicates immune-mediated mechanisms in AD^12,13^, but the causal roles for many other gene expression signatures remain unsettled.

**Fig. 1:**
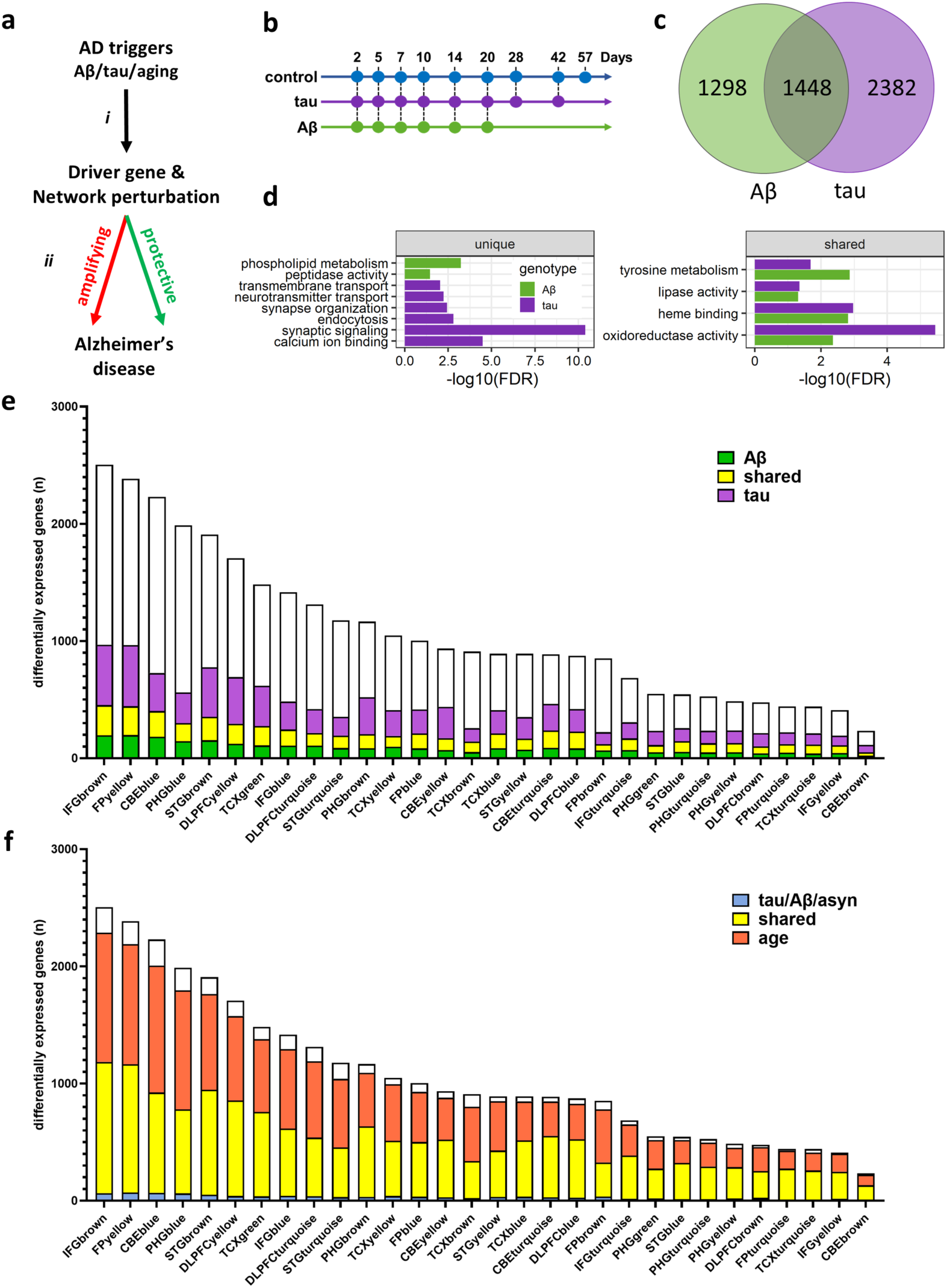
Cross-species dissection of Alzheimer’s disease (AD) pathologic triggers. **a**, Hypothetical causal chain linking AD triggers, gene expression perturbations, and Alzheimer’s disease pathophysiology. Perturbations in gene expression (up- or down-regulation) that are causal modifiers of disease pathophysiology may be either amplifying or protective. Amplifying changes are pathogenic, promoting AD pathogenesis; whereas protective changes are compensatory, attenuating AD progression. **b**, Control and AD transgenic models (n=3 replicates for each genotype) were profiled for locomotor behavior and gene expression using RNA-sequencing, including up to 9 age timepoints between 2 and 57 days. AD model genotypes were as follows: Aβ (*elav-Gal4/+; UAS-Aβ/+*) and tau (*elav-Gal4/+; UAS-tau/+*). Both wildtype (*w*^1118^) and driver (*elav-GAL4/+*) controls were evaluated. See also Extended Data Fig. 1. **c,** Venn diagram highlighting shared and unique differentially expressed genes following pan-neuronal expression of Aβ or tau. Linear regression was performed comparing *elav>Aβ* or *elav>tau* with *elav-GAL4* driver controls, including longitudinal data (days 2-28 for Aβ and days 2-42 for tau) and adjusting for age. Statistical analysis was based on a Wald test (false discovery rate (FDR) < 0.5). See also Extended Data: Fig. 1b and Table 2. **d**, Plots highlight shared and unique pathways perturbed in Aβ and tau transgenic flies, based on gene ontology term enrichment. Statistical analysis based on the hypergeometric overlap test (FDR < 0.5). **e**, Cross-species analysis highlights genes within human AD-associated coexpression modules, for which expression of conserved *Drosophila* homologs are triggered by Aβ (green), tau (purple), or both Aβ/tau (yellow). Each bar indicates the total module size, based on the total number of conserved genes. See also Extended Data Table 5. **f**, Cross-species analysis highlights genes within human coexpression modules, for which conserved *Drosophila* homologs are differentially expressed in response to AD and related dementia pathologic triggers (blue: Aβ, tau, or alpha-synuclein), aging (orange), or both aging and disease triggers (yellow). See also Extended Data Fig. 1c.

Several major challenges hinder the comprehensive functional dissection of the causal chain in AD pathogenesis. First, aging and other comorbid brain pathologies can confound interpretation of differentially-expressed genes, making it difficult to identify the specific changes induced by AD pathology, including amyloid-beta (Aβ) plaques and tau neurofibrillary tangles. Second, AD pathophysiology is a dynamic process that evolves over decades, but human brain transcriptional profiles can only be examined at one timepoint following death. Third, a major obstacle remains to pinpoint those gene expression changes that are causal, which we define as having the potential to alter AD onset, progression, and/or neurodegeneration. Among causal gene expression perturbations, it is further important to differentiate amplifying versus protective roles in disease, that is whether they promote or rather compensate for AD pathophysiology (Fig. 1a). Lastly, because changes in the transcriptome comprise correlated, modular coexpression networks with hundreds or thousands of genes per module, it is also necessary to dissect the fine-scale structure to identify key drivers with the potential to restore regulatory homeostasis.

By contrast with expression profiling of human postmortem brain tissue, animal models enable controlled experimental manipulations that isolate specific molecular triggers (e.g., Aβ vs. tau) along with longitudinal study designs that account for the dynamic impact of aging. Mice harboring *amyloid precursor protein* (*APP*) or *microtubule associated protein tau* (*MAPT*) transgenes recapitulate gene expression signatures similar to AD human brain transcriptomes, including dysregulation of immune and synaptic processes^2,14–19^. Moreover, genetic manipulations of immune regulators such as *TREM2* or *NFkB* in microglia, have been demonstrated to modify neurodegeneration, further supporting causal roles for these pathways^20,21^. However, the coexpression modules derived from the AMP-AD human brain transcriptome meta-analysis range in size from 500 to 5000 genes, creating an impediment to evaluate a larger number of potential targets. High-throughput systematic genetic dissection, or systems genetics, to pinpoint causal genes and network drivers is not feasible in mouse models, requiring alternative strategies. In the fruit fly, *Drosophila melanogaster,* pan-neuronal expression of the secreted human Aβ peptide or MAPT/tau protein triggers misfolding and aggregation similar to amyloid plaque or neurofibrillary tangle pathology, respectively, along with age-dependent, progressive neurodegeneration^22–25^. We recently showed that pan-neuronal expression of tau in flies triggers conserved gene expression signatures strongly overlapping with differentially expressed genes from AMP-AD human brain transcriptomes^26^. Here, we deploy cross-species approaches for functional dissection of AD gene expression signatures, differentiating specific and shared responses to Aβ, tau, and/or aging, and mapping the key driver genes and causal networks that mediate the impact of disease pathology on progressive central nervous system (CNS) dysfunction.

## RESULTS

### Cross-species dissection of causal pathologic triggers

Our previously published AMP-AD meta-analysis of postmortem brain RNAseq defined 3,774 AD differentially expressed genes and implicated 30 human brain gene coexpression network modules, relying on an integrated clinical-pathologic diagnosis^2^. The modules were independently derived from 7 brain regions and are organized into 5 overlapping consensus clusters (denoted A-E; Extended Data Table 1). To address potential molecular specificity (arrow *i* in Fig. 1a), we leveraged available complementary differential gene expression analyses^27^ based on Aβ or tau neuropathology and examined enrichment across all modules (Extended Data Table 1). All 30 modules were associated with AD pathology, including both predominantly up- and down-regulated modules. As expected, since amyloid and tau pathology co-occur and are highly correlated in AD, most modules showed consistent enrichment with differentially expressed genes from both analyses. Next, to determine which transcriptional changes might be directly caused by AD pathology and to further differentiate Aβ versus tau as potential triggers we turned to *Drosophila* transgenic models. As introduced above, pan-neuronal expression of either the secreted human 42-amino-acid Aβ peptide or the tau protein (wild-type 2N4R isoform) recapitulates age-dependent neuronal loss and progressive CNS dysfunction, including locomotor impairment (Extended Data Fig. 1a)^25^. We used previously published *Drosophila* strains^28,29^ harboring human transgenes responsive to the yeast Upstream Activating Sequence (UAS); whereby, expression is directed selectively to neurons by the *Elav-GAL4* driver^30^. *Elav>Aβ*, *Elav>tau*, or control flies (*Elav-GAL4 /+* or *w*^1118^) were aged, and longitudinal transcriptome profiles were generated at up to 9 time points (Fig. 1b). We used multivariate linear regression to model the impact of each AD pathologic trigger on gene expression, including age as a covariate. Aβ and tau induced 2,746 and 3,830 differentially expressed genes, respectively (Fig. 1c, Extended Data: Fig. 1b and Table 2). There was substantial overlap in the transcriptional signature induced by Aβ or tau, but specific changes were also identified. For example, based on enrichment for gene ontology terms, genes related to lipid and oxidative metabolism were perturbed in both models (Fig. 1d and Extended Data Table 3). Conversely, regulators of synaptic transmission and calcium related processes were specifically enriched among tau-induced, differentially expressed genes, whereas glycerophospholipid metabolism was specifically highlighted in the Aβ transgenic model. Similar functional enrichments also characterized several of the AD-associated human coexpression modules from the AMP-AD meta-analysis (Table 1). For example, module PHGbrown which is down-regulated in AD is characterized by abundant neuronal and synaptic genes^2^.

**Table 1.**
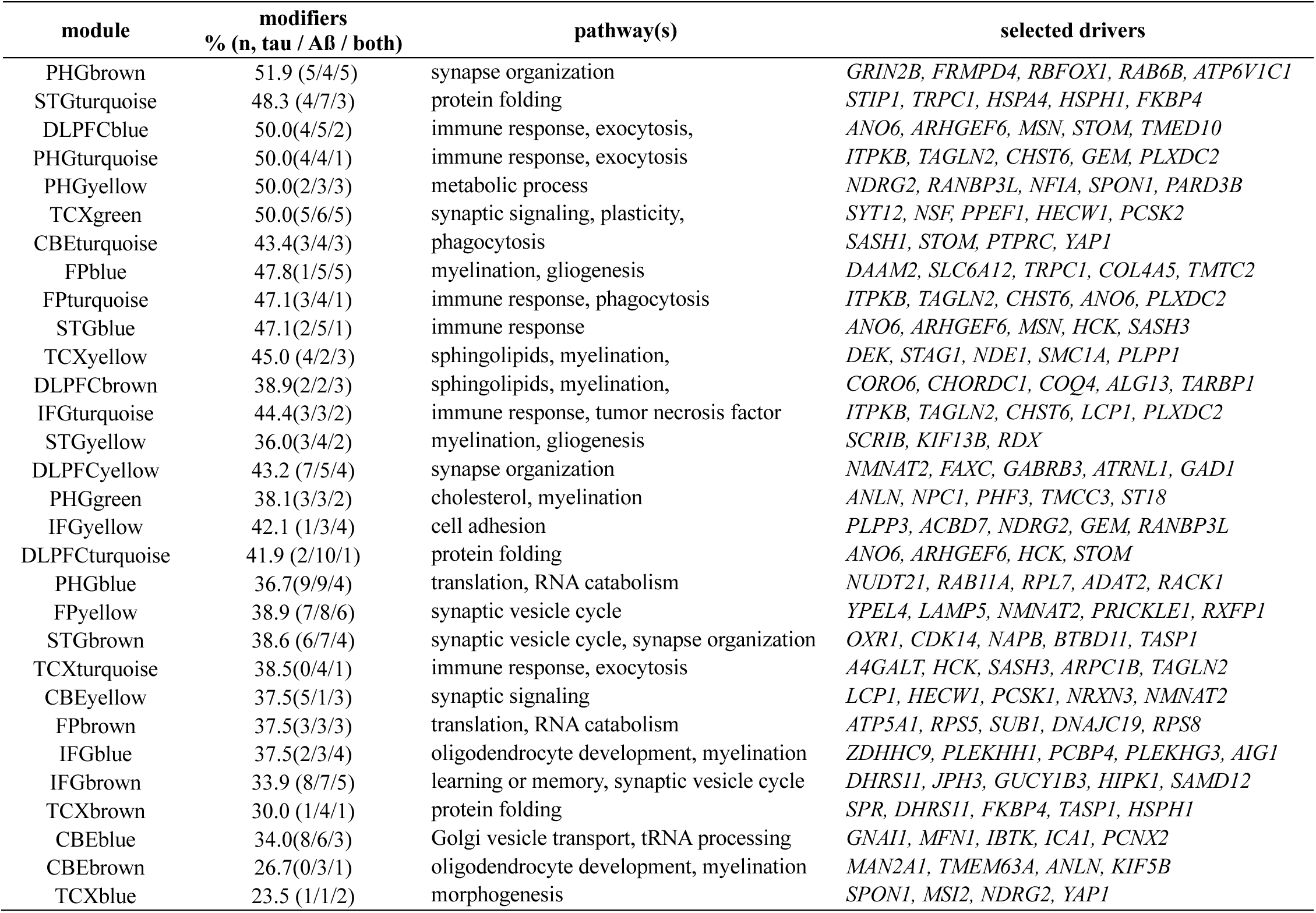
AMP-AD coexpression modules and cross-species screen results.

To more directly integrate the AMP-AD human consensus coexpression modules with *Drosophila* differential expression data, we mapped gene homologs across species (Extended Data Table 4). Most human coexpression modules (24 out of 30) were significantly enriched for fly gene homologs showing differential expression in response to tau (n=25) and/or Aβ (n=12) (Fig. 1e and Extended Data Table 5). Based on cross-species overlaps, we infer that most modules are at least partially activated by both AD triggers. We therefore partitioned each module into discrete Aβ- and/or tau-responsive gene sets, along with a shared component due to the substantial transcriptome overlap between the Aβ and tau transgenic flies (Fig. 1e). Overall, among all conserved genes within each module, the proportion of genes responsive to AD pathologic triggers varied (range_Aβ_ = 14-22%; range_tau_ = 18-43%). For example, among 887 conserved genes in CBEturquoise, which is related to phagocytosis and enriched for microglial markers, 279 (31.5%) or 455 (51.3%) genes, respectively, significantly overlapped with Aβ- or tau-induced differentially expressed genes from transgenic flies, and these overlaps were highly unlikely to have occurred by chance (p_Aß_ = 4.1×10^−6^; p_tau_ = 1.3×10^−16^). In other cases, such as for the PHGbrown module, we note a more selective enrichment for differentially expressed genes from the *Drosophila* tau model (p = 1.2×10^−7^), but not Aß (p = 0.74).

In human brains, amyloid plaques and tau tangles frequently co-occur with other pathologies, such as alpha-synuclein (⍺Syn) Lewy bodies, which can increase vulnerability for cognitive impairment in AD and also cause Lewy body dementia^31^. Pan-neuronal expression of human ⍺Syn (*Elav>⍺Syn*) in *Drosophila* causes Lewy-body like protein aggregation, dopaminergic and other neuron loss, and progressive locomotor impairment^32–34^. We therefore obtained complementary transcriptome profiles from ⍺Syn transgenic flies using identical aging timepoints and experimental conditions, facilitating comparisons with the complementary results from *Elav>tau* and *Elav>Aβ* models. Overall, there was substantial overlap in brain gene expression changes triggered by distinct disease models, with 875 common DEGs among the 3 genotypes and 38-49% of differentially expressed genes being shared among pairwise comparisons of perturbations triggered by Aβ, tau, or ⍺Syn (Extended Data: Fig. 1b and Table 2). Since aging has a strong impact on the brain independent of disease pathologies, we also defined aging-induced gene expression changes from control flies (*w*^1118^). Consistent with our prior work^26^, aging had a profound impact on the transcriptome, significantly altering expression of 10,717 genes. Moreover, approximately 83% of expression changes observed in the neurodegenerative models were also perturbed in aging control animals (Extended Data Fig. 1b). We therefore extended the cross-species approach to pinpoint putative ⍺Syn- and aging-induced differentially expressed genes among the human AD-associated coexpression modules (Fig. 1f and Extended Data Fig. 1c). In flies, ⍺Syn altered the expression of 24-39% conserved genes from each human coexpression module, whereas aging had an even greater effect (86-95% genes). Overall, the aggregate impact of aging along with Aβ, tau, and ⍺Syn pathologies potentially explains the majority of human brain gene expression changes of AD (Fig. 1f).

### Cross-species dissection of brain expression networks and behavior

The preceding analyses highlight extensive conservation between the transcriptional changes observed in human brain and *Drosophila* models of AD, other AD-related dementias, and aging. We next examined if this conservation extends to the network level. We independently performed weighted gene coexpression network analysis (WGCNA)^35^ using transcriptome profiles from the *Drosophila* adult brain. For optimal power, we leveraged all available *Drosophila* transcriptome data, including longitudinal profiles obtained from control, Aβ, tau, or ⍺Syn flies. The combined dataset comprises 147 total samples, including bulk RNAseq in triplicate from fly heads of 7 distinct genotypes and up to 9 timepoints between 2 and 57 days of aging. Importantly, the network detection algorithm is implemented independent of sample age and genotype in order to identify gene sets that covary across a range of conditions. Overall, WGCNA identified 19 modules ranging from 32 to 4277 genes (Extended Data: Fig. 2 and Table 2). All 30 AMP-AD human coexpression modules significantly overlap with at least one *Drosophila* coexpression module (Fig. 2a and Extended Data Table 6). Several overlapping fly and human modules also showed similar enrichment for cell-type specific markers and biological pathways (Extended Data: Fig. 2b and Table 7). For example, *Drosophila* module fM1 is strongly enriched for neuronal markers and genes that mediate synaptic signaling (GO:0099536, p<1.0×10^−15^) and overlaps significantly with PHGbrown (p=1.7×10^−50^). Overall, these results support conservation of gene regulatory networks present in the adult *Drosophila* brain, including models of AD and related neurodegenerative disorders.

**Fig. 2:**
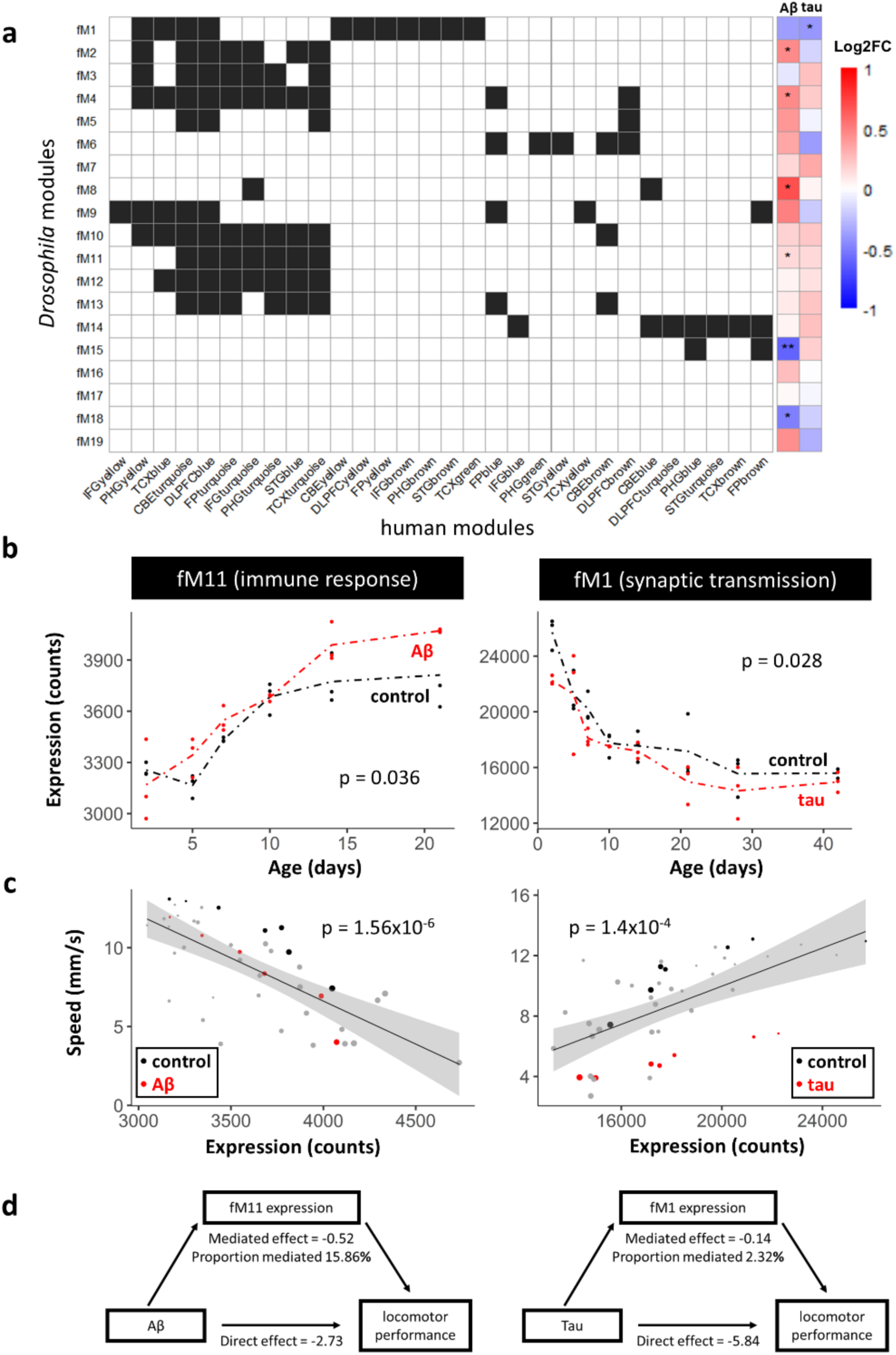
Cross-species dissection of brain gene expression networks and behavior. **a**, Overlap between *Drosophila* WGCNA modules (rows) and AMP-AD human gene coexpression modules (columns). Black shading indicates significant overlaps based on the hypergeometric overlap test (false discovery rate < 0.05). The heatmap, at right, indicates the direction and magnitude of mean differential expression for module genes in either the Aβ (*elav-Gal4/+; UAS-Aβ/+*) or tau (*elav-Gal4/+; UAS-tau/+*) *Drosophila* models when compared with controls (*elav-GAL4/+*), using linear regression models including longitudinal data (days 2-28 for Aβ and days 2-42 for tau) and adjusting for age. Statistical analysis based on the likelihood ratio test applied with significance-level adjusted based on false discovery rate (*, p<0.05, **, p<0.01). See also Extended Data: Fig. 2-4 and Tables 6, 8. **b**, The conserved *Drosophila* coexpression modules fM11 and fM1 are significantly up- and down-regulated, respectively, in response to Aβ and tau. Mean expression for module genes are shown for AD transgenic models (red, *elav>Aβ* or *elav>tau*) or *elav-GAL4* driver controls (black). See also Extended Data: Fig. 3,4 and Table 8. **c**, Scatter plots demonstrate a strong relation between fM11/fM1 mean module expression and locomotor behavior (climbing speed), based on Pearson correlation analysis (significance adjusted based on false discovery rate). The plots include data from 44 different experimental conditions, including 7 genotypes and evaluated at 7 aging timepoints between 2 and 28 days. The color key indicates the relevant AD transgenic model (red) and controls (black), with all other conditions noted in grey, and age denoted by size of the data point. Gray shading denotes the 95% confidence interval (standard error of the mean) for the line of best fit. See also Extended Data Fig. 5. **d**, Mean expression of fM11 and fM1 genes partially mediate the impact Aβ and tau, respectively, on progressive CNS dysfunction (locomotor behavior). Mediation analysis considered the association between Aβ or tau and locomotor performance phenotype, decomposing into the component explained by module expression, and the residual, unexplained component (direct effect). Proportion mediated indicates the fraction of the mediated effect to the total effect.

We hypothesize a causal chain in which disease triggers, such as Aβ, tau, and aging, modulate brain gene expression networks, which subsequently impact CNS function (Fig. 1a). In nested linear regression models considering the joint impact of AD pathologic triggers and aging, Aβ significantly perturbed 6 coexpression modules, whereas tau altered expression of 1 module (Fig. 2a-b and Extended Data: Figs. 3,4 and Table 8). Next, in order to directly model the link between brain transcriptional networks and CNS function/behavior, we leveraged locomotor performance data (negative geotaxis) generated from the same fly cohort used for RNA-sequencing. Overall, mean expression of 6 out of 19 transcriptional modules were significantly correlated with locomotor performance (Fig. 2c and Extended Data Fig. 5), including networks enriched for genes that regulate the immune response (fM11), synaptic transmission (fM1), and endocytosis/vesicle trafficking (fM8) (Extended Data Table 7). Lastly, using multivariate mediation models^36^, we asked whether these module expression levels explain the impact of Aβ and/or tau on progressive CNS dysfunction (Fig. 2d). Among our results, Aβ triggered up-regulation of module fM11, which in turn strongly mediated the association of Aβ with locomotor impairment (model beta reduced by 16%) (Fig. 2d, left). Interestingly, fM11 overlapped significantly with STGblue and other AMP-AD microglial modules and is similarly enriched for immune pathways. Complementary analyses indicate more modest attenuation of tau-induced CNS dysfunction. For example, tau induces down-regulation of module fM1, which in turn, partially mediates the tau association with locomotor impairment (model beta reduced by 2%) (Fig. 2d, right). In sum, these cross-species results—integrating *Drosophila* AD models, behavioral assays, and brain transcriptome profiles—highlight conserved gene regulatory networks that respond to neurodegenerative pathology and may mediate the downstream consequences for progressive behavioral impairment.

**Fig. 3:**
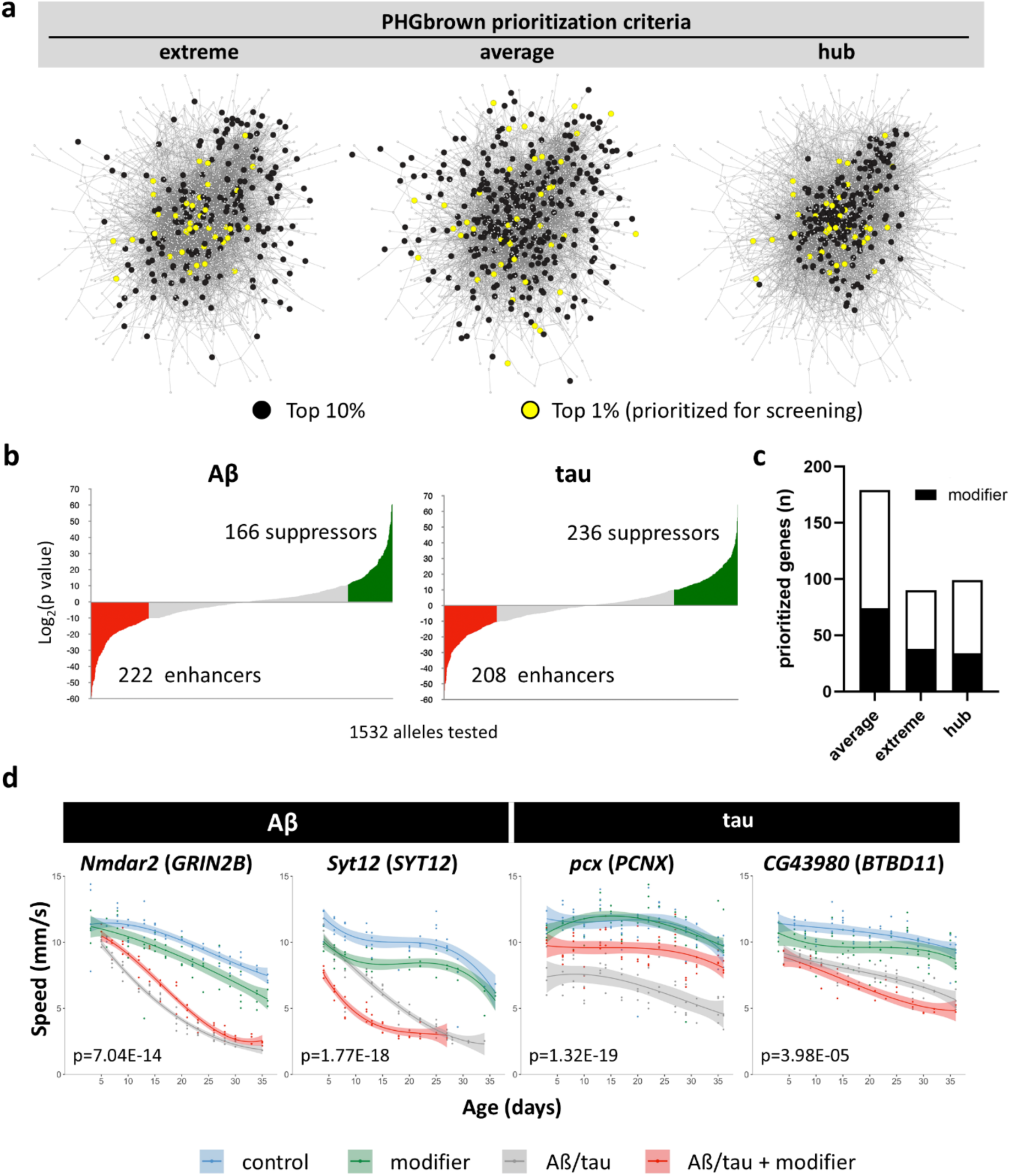
Prioritization and screening for causal drivers from AD brain expression networks. **a**, Schematic network plot for the PHGbrown coexpression module, showing 3 independent and non-overlapping criteria for selection of candidate drivers for experimental manipulations. The top-ranked 10% (black) or 1% (yellow) gene are highlighted for each prioritization criterion (extreme, average, hub). See also Extended Data Table 9. **b**, Plot shows summary statistics for results of all 1,532 alleles tested, denoting the number of recovered enhancers (red) and suppressors (green) of Aβ- and/or tau-mediated locomotor behavior impairment. Each modifier test included at least n > 4 replicates of 10 animals each examined at 6 or more timepoints over 28 days of aging. Statistical comparisons based on one-way ANOVA considering three nested models (genotype, genotype + time, and genotype*time) and applying a significance threshold of p<0.05, following Holm-Bonferroni adjustment. See also Extended Data Table 11. **c**, All three prioritization criteria identify causal modifiers of Aβ- and/or tau-mediated CNS dysfunction. Driver genes were based on consistent, statistically significant modifier effects from at least 2 independent allele strains. **d**, Representative results from the screen of prioritized PHGbrown candidate drivers in the Aβ (*elav-Gal4/+; UAS-Aβ42/+*) and tau (*elav-Gal4/+; UAS-tau/+*) transgenic fly models. Plots show the results of longitudinal locomotor behavior assays (climbing speed during startle-induced negative geotaxis). The *Drosophila* gene homolog tested is noted above each plot, along with human gene in parentheses. The following RNAi knockdown or loss-of-function alleles were tested in heterozygosity: *Nmdar2(NIG14794R-3III); Syt12(f00785)*; *pcx*(*v29893);* and *CG43980(v107170)*. Natural spline curve with 3 degrees of freedom was fit to data, with shading to denote the 95% confidence band. See also Extended Data: Fig. 6 and Table 11.

**Fig. 4:**
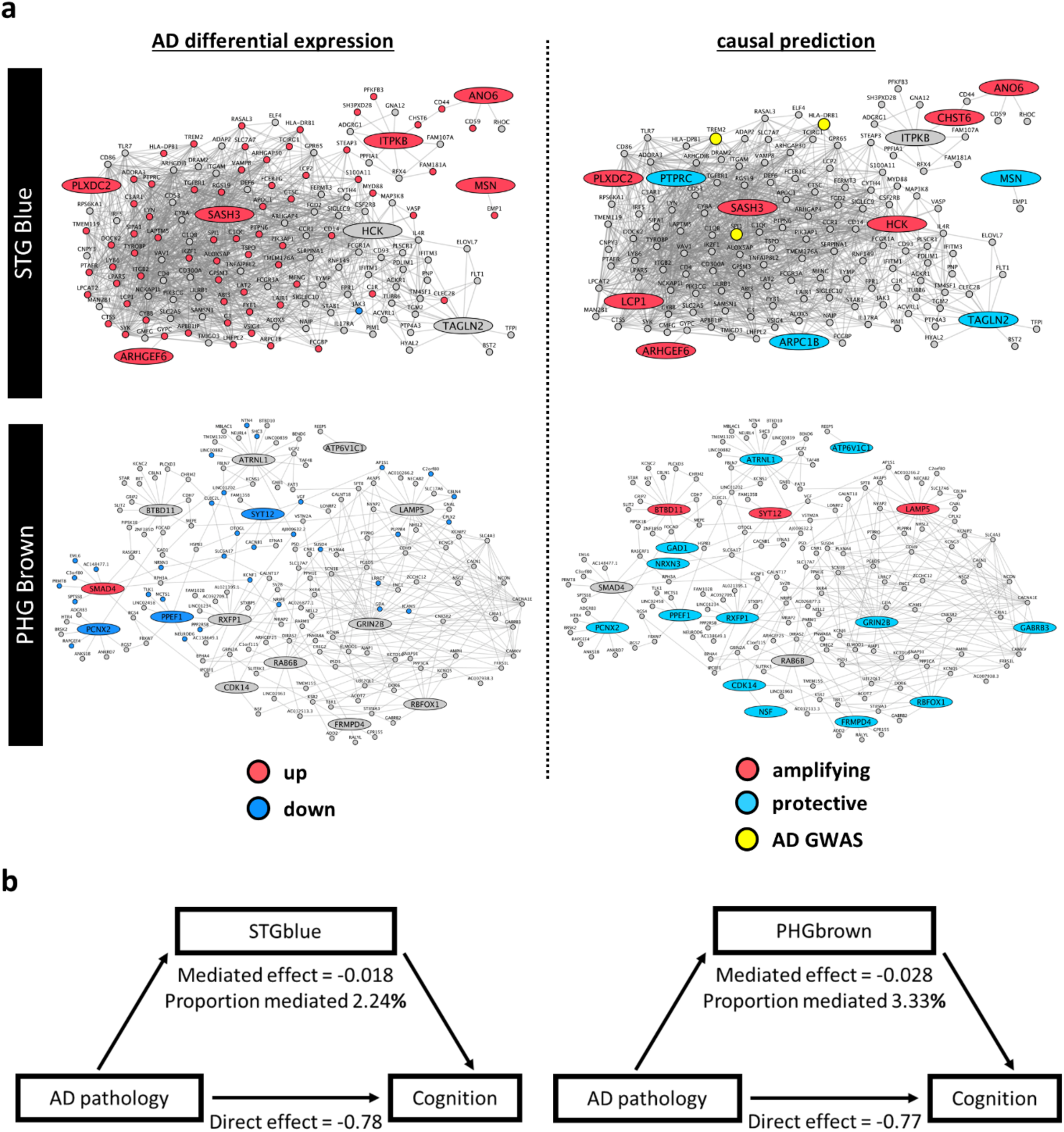
Alzheimer’s disease (AD) causal gene expression networks. **a**, Differential AD-associated gene expression and causal driver predictions are shown for the STGblue and PHGbrown networks. In order to derive subnetworks graphs, we plotted the human homologs of network modifier genes (large oval nodes) along with all first-order neighbors (small circle nodes). Edges indicate coexpression relationships between genes. Left: Up- or down-regulated expression is indicated using red or blue, respectively, based on the published AMP-AD case/control brain transcriptome meta-analysis. Nodes colored grey represent genes which were not significantly differentially expressed between AD and control samples in the AMP-AD data. Right: Integrating results from the *Drosophila* modifier screen (enhancers or suppressors of Aβ/tau-induced locomotor impairment), we inferred whether causal drivers are predicted to be amplifying or compensatory (red or blue, respectively). Amplifying gene expression perturbations are expected to promote disease pathophysiology, whereas compensatory changes are predicted to be protective. Genes that localize to established AD risk loci are highlighted as yellow nodes. Genetic manipulations of fly homologs of ITPKB from STGblue, and both *SMAD4* and *RAB6B* from PHGbrown were compatible with bi-directional effects (colored gray). See also Extended Data: Fig. 7 and Table 12. **b**, The mean expression of STGblue and PHGbrown genes partially mediate the impact of AD pathology on cognitive performance, based on analyses of clinical-pathologic data from the Religious Orders Study and Rush Memory and Aging Project (n=636, mean age at death=88.7, age range=67-108, female sex=64%, AD=40.1% non-demented=58%). Mediation analysis considered the association between global AD pathology (quantitative assessments of amyloid plaques and tau neurofibrillary tangles) and global cognitive performance in evaluations proximate to death (summary measure from 19 cognitive tests), decomposing into the component explained by module expression (average causal mediation effect), and the residual, unexplained component (average direct effect). Proportion mediated indicates the fraction of the mediated effect to the total effect. Linear regression models relating pathology, cognition, and gene expression included covariates for age at death, postmortem interval and sex. Significance testing from 1000 simulations with the Benjamini-Hochberg false discovery rate (mediation p_STGblue_ = 0.048; p_PHGbrown_ = 0.036).

### Prioritizing network drivers

Rather than a collection of co-equal members, we hypothesize that AD-associated coexpression modules possess a fine-scale, hierarchical structure, in which selected gene drivers regulate the broader network and may subserve causal roles in disease^37^. We therefore sought to systematically prioritize candidate driver genes within each regulatory network. We considered three distinct and complementary bioinformatic criteria for ranking genes within each module (Fig. 3a). Since the 30 human AMP-AD modules were identified based on their enrichment for AD differentially expressed genes we first considered the magnitude of differential expression, prioritizing genes within networks with the most extreme expression differences between AD cases and controls. Second, we ranked genes based on correlation with the first principal component of expression for each module, highlighting network “eigengenes” that best approximate the average module transcriptional activity. Lastly, we identified hub genes based on network connectivity, or degree, reflecting the number of edges due to pairwise coexpression of module gene members. Overall, the 3 selected parameters (extreme, average, and hub) define largely independent and non-overlapping criteria for ranking candidate key drivers within each module (Fig. 3a and Extended Data Table 9). Further, the nominated driver genes recapitulate the cell-type specific and functional signatures of each module (Extended Data Table 10). For example, the 614 prioritized drivers from PHGbrown, representing the top 10% genes ranked using each criteria, were strongly enriched for synaptic signaling gene ontology terms (GO:0099536, p=5×10^−85^) similar to the entire module (n=2,123 total genes). Consistent results were also obtained when separately considering PHGbrown drivers nominated based on each of the distinct prioritization criteria (p_average_=2.5×10^−33^; p_extreme_=1.3×10^−6^; p_hub_=1.2×10^−54^).

### Screening for causal drivers of Aβ- and tau-mediated neurodegeneration

Our preceding analyses identify conserved transcriptional modules as potential mediators of Aβ- or tau-triggered neurodegeneration, along with promising candidate key drivers within these networks. Nevertheless, without directed experimental manipulations, it is challenging to definitively establish whether differentially expressed genes identify upstream causal factors versus downstream, reactive changes. We therefore next applied our cross-species strategy to determine whether prioritized driver genes and modules are causal (arrow *ii* in Fig. 1a), screening for *in vivo* modifiers of Aβ- and/or tau-induced progressive neuronal dysfunction. We focused on gene candidates representing the top 1% ranked by each driver criterion, calibrating our selections based on module size and overlaps (see Methods). Notably, genes prioritized based on extremes of differential expression were less conserved (55.7%), when compared with those selected based on average expression (66.1%) or hub-like connectivity (68.3%). Overall, we prioritized 357 conserved candidate AMP-AD driver genes, including 90 extreme, 179 average, and 99 hub-like genes (Extended Data Table 9). Our screen thus considered an average of 28 (range=13-61) conserved candidate driver genes per module. Because of overlaps, a larger number of genes were in fact tested per module (mean=67, range=28-118), whereby predicted drivers from one module may be members of a related module from a different brain region. In cases where a human driver gene candidate had multiple potential homologs, we considered up to 3 well-conserved candidates based on the Drosophila RNAi Screening Center Integrative Ortholog Prediction Tool (DIOPT)^38^. Thus, for the 357 human candidate genes, our screen evaluated a total of 433 *Drosophila* homologs. We obtained 1,532 genetic strains, including RNA-interference (RNAi) transgenic lines and classical loss-of-function alleles (average=3.5 independent lines per target gene, range=2-8), as well as lines predicted to activate target gene expression. In 115 out of 433 fly genes evaluated, the availability of both loss- and gain-of-function alleles permitted reciprocal tests of the consequences for both down- and up-regulation of candidate drivers. All lines were tested by crossing with *Drosophila* strains in which secreted Aβ or wild-type 2N4R tau are expressed throughout the nervous system, using the identical *Elav>Aβ* and *Elav>tau* models characterized above. We utilized an automated locomotor behavioral assay which is amenable for high-throughput genetic screening^39^. The climbing speed of adult flies was evaluated longitudinally between 1 and 3 weeks of age, scoring for enhancement or suppression of the locomotor phenotypes caused by Aβ- or tau-induced neuronal dysfunction. RNAi knockdown was activated pan-neuronally via the *Elav-GAL4* driver, which also directs expression of Aβ/tau transgenes. All alleles were tested in heterozygosity. While *Drosophila* RNAi lines are designed for optimal specificity^40,41^, we only considered genes as modifiers when supported by consistent evidence from at least 2 independent RNAi strains or other alleles, thereby further minimizing the possibility of off-target or genetic background effects.

Overall, our primary screen identified a total of 134 *Drosophila* genetic modifiers, homologous to 141 predicted human gene drivers (Fig. 3b, Table 1, and Extended Data Tables 9, 11). All three prioritization criteria led to the successful identification of causal modifiers of Aβ- and tau-induced progressive neuronal dysfunction. Proportionally, candidate drivers prioritized based on either extreme or average differential expression were somewhat more likely to be validated as modifiers (42.2 and 41.3%, respectively) than the hub-like genes (34.3%) (Fig. 3c). We identified similar numbers of Aβ (n=93) and tau (n=88) modifiers, including 41 genes that interact with both AD models following knockdown and/or activation. The screen highlights an average of 11 modifiers per module (range = 4 to 22 modifiers), pinpointing those regulatory systems comparatively enriched (or depleted) for candidate drivers with causal evidence (Table 1). PHGbrown had the highest overall modifier hit rate (52%); whereas TCXblue, enriched for genes with roles in developmental morphogenesis, had the lowest number of modifiers recovered from our screen (∼24%). While most modules identified modifiers of both fly models, others showed potential specificity. For examples, DLPFCturquoise is characterized by disproportionate numbers of Aβ modifiers (n=10) versus tau (n=2). Table 1 summarizes tallies of all recovered modifiers from each module, notable representative driver genes, and implicated pathways.

All enhancer strains (371 RNAi and other alleles) were retested to examine the consequences for locomotor behavior in the absence of Aβ or tau (e.g., *elav>RNAi / +*). Based on these results, we further classified all enhancer alleles as either additive or synergistic modifiers (Extended Data Table 11). Additive modifiers, in which genetic manipulations cause locomotor independent of Aβ or tau, are excellent candidates to mediate alternate biological pathways important for neurodegeneration, such as requirements for neuronal maintenance with aging. We identified 39 *Drosophila* genes with modifier alleles that enhanced Aβ or tau additively. Interestingly, module STGyellow, which is characterized by oligodendrocyte marker genes, was particularly enriched in causal drivers for which fly homologs showed CNS requirements independent of Aβ or tau (5 out of 25 genes evaluated), including *MYRF, SCRIB, KIF13B, ST18,* and *PLPP1*.

### Defining and validating human AD causal subnetworks

The AMP-AD consensus coexpression modules are large, ranging from 480 to 4519 gene members; therefore, our genetic screen examined a systematically prioritized subset of candidate driver genes. To define putative causal networks, partition subnetworks, and further define potential mechanisms, we next integrated our *Drosophila* genetic screen results with human network structure (Fig. 4a and Extended Data 7). Starting with the experimentally validated driver genes as seeds, we plotted all first degree neighbors from the consensus coexpression network and also determined whether each driver and other member genes are up- or down-regulated in human postmortem brain tissue, based on the published AD case/control differential expression meta-analysis^2^. We next summarized whether the AD-associated perturbation of each validated driver gene (e.g., increased or decreased differential expression) is predicted to either promote or protect against disease pathogenesis (Fig. 1a), based on the corresponding results from our genetic manipulations of the *Drosophila* gene homolog in Aβ/tau models (Extended Data Table 12). Gene expression changes inferred to aggravate Aβ- and/or tau-mediated neurotoxicity were classified as disease amplifying/pathogenic gene expression changes. By contrast, differentially expressed genes inferred to ameliorate Aβ- and/or tau-induced brain injury were classified as compensatory/protective genes. Together, these results define a collection of 30 AD causal subnetworks based on robust meta-analyses conducted in large human autopsy studies and integrated with results from systematic genetic manipulations to validate likely causal drivers (Fig. 4 and Extended Data: Fig. 7 and Tables 12).

We next systematically examined whether any of the causal subnetworks identify genes associated with AD risk (Extended Data Table 13). Leveraging publicly available summary statistics from 2 AD genome-wide association studies (GWAS)^42,43^, we deployed the multi-marker analysis of genomic annotation (MAGMA)^44^ tool, to examine whether each subnetwork is enriched for AD susceptibility variants.

MAGMA computes an overall gene-set test statistic, including adjustments for gene size and regional linkage disequilibrium. Two subnetworks, STGblue and PHGturquoise, were consistently enriched for AD susceptibility variants (p<0.05). Similar results were not observed in analyses of the full module, nor in subnetworks generated from all prioritized genes (Extended Data Table 16); therefore, integration of the *Drosophila* experimental results enhances power to detect causal networks. Moreover, we did not detect similar enrichment when using GWAS summary statistics from for either Parkinson’s disease^45^ or height^46^, consistent with specificity for AD risk.

The STGblue and PHGturquoise causal subnetworks are similarly enriched for microglial expression signatures (p_STGblue_=1.4×10^−4^; p_PHGturquoise_=6.4×10^−6^) and genes that regulate innate immune mechanisms (GO:0006955, p_STGblue_= 2.2×10^−32^; p_PHGturquoise_= 8.7×10^−29^) (Extended Data Table 14). Indeed, AD GWAS meta-analysis results are strongly enriched for microglial and innate immune pathways^13^ and these subnetworks are notable for including genes from AD susceptibility loci. For example, the STGblue subnetwork includes *TREM2*, *HLA-DRB1*, and *SPI1*, which all have roles in immune modulation (marked yellow in Fig. 4). Interestingly, although these genes are poorly conserved in *Drosophila*, they are coexpressed with conserved, prioritized driver genes that were validated as modifiers in our screen. Genes in the STGblue subnetwork are predominantly up-regulated in AD^2^. Among the 8 STGblue modifier genes, we can infer 5 amplifying and 3 compensatory genes. For example, *SASH3* is an STGblue hub driver candidate, encoding an immune signaling adapter protein^47^. Loss-of-function of the fly gene homolog *SKIP*, either RNAi-mediated knockdown or heterozygosity for an insertional allele, suppresses Aβ and tau-mediated locomotor impairment (Extended Data Table 11). We therefore infer that up-regulation of *SASH3* along with 4 other STGblue drivers may promote AD pathogenesis. Since most of our tested alleles were RNAi transgenes under the control of the pan-neuronal *elav-GAL4* driver, our results are highly suggestive of cell autonomous, causal roles within neurons, besides the well-established microglial role of these genes in innate immunity. Indeed, prior studies in flies, mice, and human brains support the expression of immune regulators and effectors in many neuronal subtypes^48^

Among our screen results, PHGbrown had the highest hit-rate, with 14 out of 27 candidate driver genes modifying Aβ (n=9) or tau (n=10) neurotoxicity, with manipulations of 5 genes modifying both AD triggers (Fig. 4 and Extended Data Table 12). The resulting 167-gene PHGbrown causal subnetwork is significantly enriched for genes that regulate synaptic function (GO:0099536, p=4.0×10^−11^) and glutamatergic neurotransmission (KEGG:04724, p=2.5×10^−4^) and is a predominantly down-regulated module in AD bulk brain tissue (Fig. 4a and Extended Data: Fig. 8 and Table 14). While the PHGbrown subnetwork was not directly associated with AD risk from our MAGMA analyses, many other endocytic genes have been identified at AD susceptibility loci, including several regulators of synaptic vesicle recycling^43^. Integrating PHGbrown differential expression with the results from screening, we infer 13 compensatory and 3 amplifying gene expression changes. For example, *GRIN2B* encoding a subunit of the N-Methyl-D-Aspartate (NMDA) glutamate receptor, was prioritized for evaluation based on its ranking as a top, hub-like driver gene candidate. RNAi knockdown of *Nmdar2*, the conserved, *Drosophila* ortholog of *GRIN2B*, suppresses Aβ-mediated locomotor impairment, consistent with a possible compensatory role. Eleven other modifier genes show similar relationships, several of which are also implicated in neurotransmission (e.g., *FRMPD4/CG42788*, *RBFOX1/Rbfox1*, *ATP6V1C1/Vha44*)^49,50^ We further confirmed that many of these genes show consistent interactions with Aβ and tau using an independent, histologic assay for age-dependent neurodegeneration in the adult *Drosophila* brain (Fig. 5 and Extended Data Fig. 9).

**Fig. 5:**
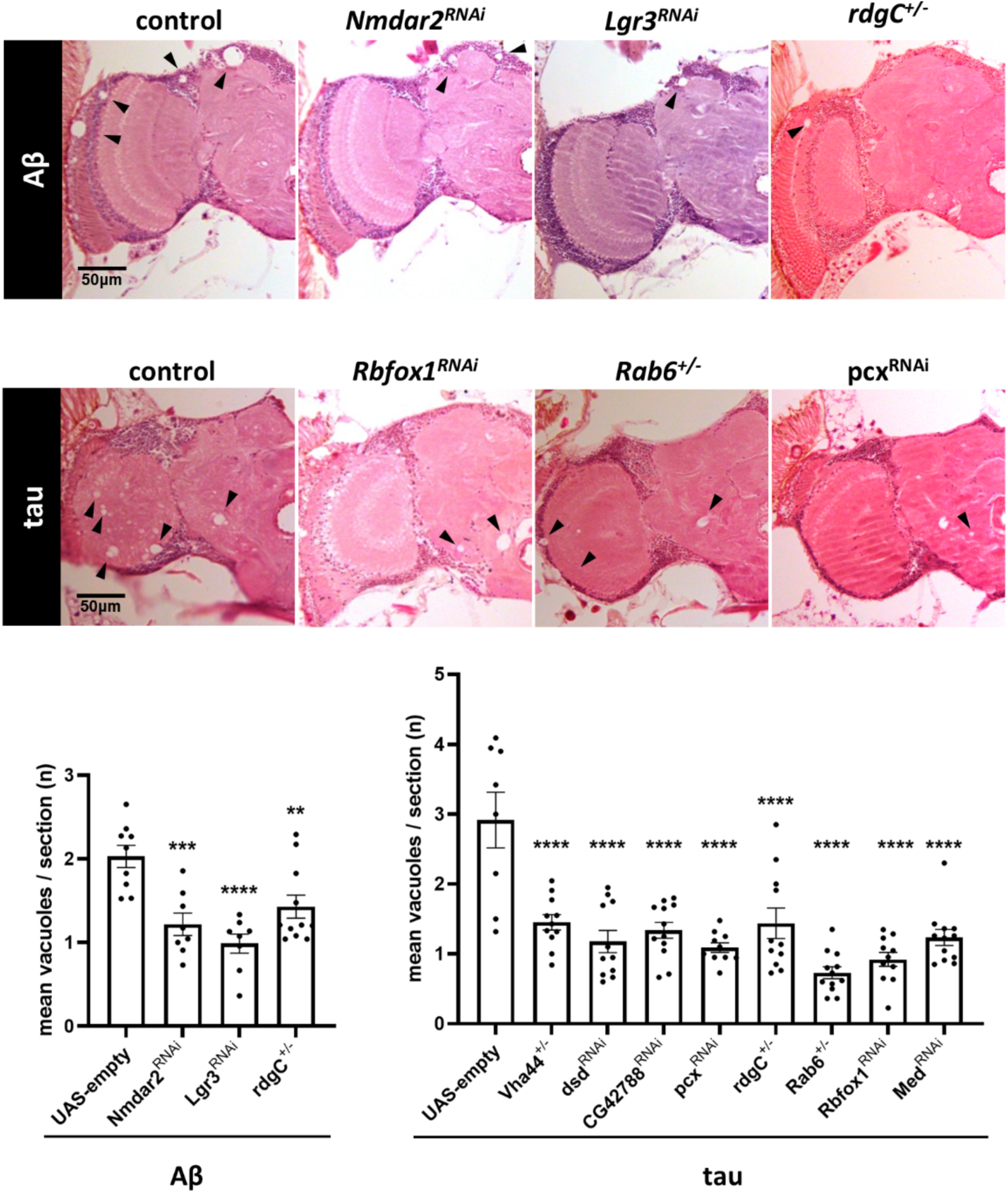
PHGbrown causal drivers modulate adult brain degeneration in *Drosophila* models. Homologs of PHGbrown causal drivers consistently modify Aβ- (*elav-Gal4/+; UAS-Aβ42/+*) or tau-(*elav-Gal4/+; UAS-tau/+*) induced neurodegeneration using an independent assay for vacuolar histopathology in the adult brain. The following RNAi knockdown or loss-of-function alleles were tested in heterozygosity: *Nmdar2(NIG14794R-3III); Lgr3(v330603)*; *rdgC(306)*; *dsd(v1106)*; *Rab6(08323)*; *CG42788(v45034)*; *Rbfox1(NIG32062Ra-3)*; *Med(v19688)*; *Vha44(MI02871)*; and *pcx*(*v29893)*. Vacuoles (arrowheads) were quantified in at least n = 8 animals at 10 or 15 days of age for tau and Aβ, respectively. Statistical analysis is based on one-way ANOVA followed by Dunnett’s test for multiple hypothesis correction. Each data point represents an individual biological replicate sample. Error bars denote the standard error of the mean. **, p < 0.01; ***, p < 0.001 **** p < 0.0001. See also Extended Data Fig. 9.

We next leveraged clinical, pathologic, and RNAseq data from human postmortem brain tissue (n=636 autopsies) in order to model the hypothetical causal chain between AD pathology, putative transcriptional networks, and downstream cognitive manifestations (Fig. 4b)^51,52^. We focused these analyses on STGblue and PHGbrown, for which our cross-species strategy highlighted potential disease-amplifying versus protective causal roles, respectively, in AD pathogenesis. Our models incorporated a quantitative summary measure of AD neuropathologic burden as an upstream trigger along with global cognitive performance from assessments proximate to death as the outcome trait. Consistent with our mediation analysis, above, considering the *Drosophila* fM1 and fM11 modules (Fig. 2c), mean expression of either the homologous human PHGbrown or STGblue explained up to 9% of the variance in cognitive impairment attributable to AD pathologic triggers. These results are consistent with a model in which AD brain pathology alters brain transcriptional programs that in turn influence downstream cognitive manifestations of disease.

### PHGbrown modulates neuronal hyperexcitability and resulting degeneration

A preponderance of evidence suggests that Aβ peptides may boost glutamatergic neurotransmission, leading to aberrant Ca^2+^ influx, CNS hyperexcitation, tau hyperphosphorylation, and ultimately neurodegeneration^53–57^. Based on our findings, we reasoned that PHGbrown may represent a conserved transcriptional response triggered by hyperexcitability that modulates brain hyperactivation injury in AD. To test whether the *Elav>Aβ* transgenic flies manifest central nervous system hyperexcitation, similar to *APP* transgenic mouse models^58^, we deployed the Transcriptional Reporter of Intracellular Ca2+ (TRIC) system^59^, in which activity-dependent calcium influx drives expression of a green fluorescent protein (GFP) reporter. Indeed, we detected a striking increase in GFP signal in the brains of 10-day-old *Elav>Aβ* animals (Fig. 6a). Next, in order to assess whether excitability mediates Aβ-induced neuronal injury, we experimentally manipulated *VGlut*, encoding the vesicular glutamate transporter, which is required for loading glutamate into synaptic vesicles. Indeed, introducing one copy of a strong *VGlut* loss-of-function allele or RNAi-mediated knockdown reduced vacuolar degeneration in *Elav>Aβ* flies (Fig. 6b and Extended Data Fig. 10a).

**Fig. 6:**
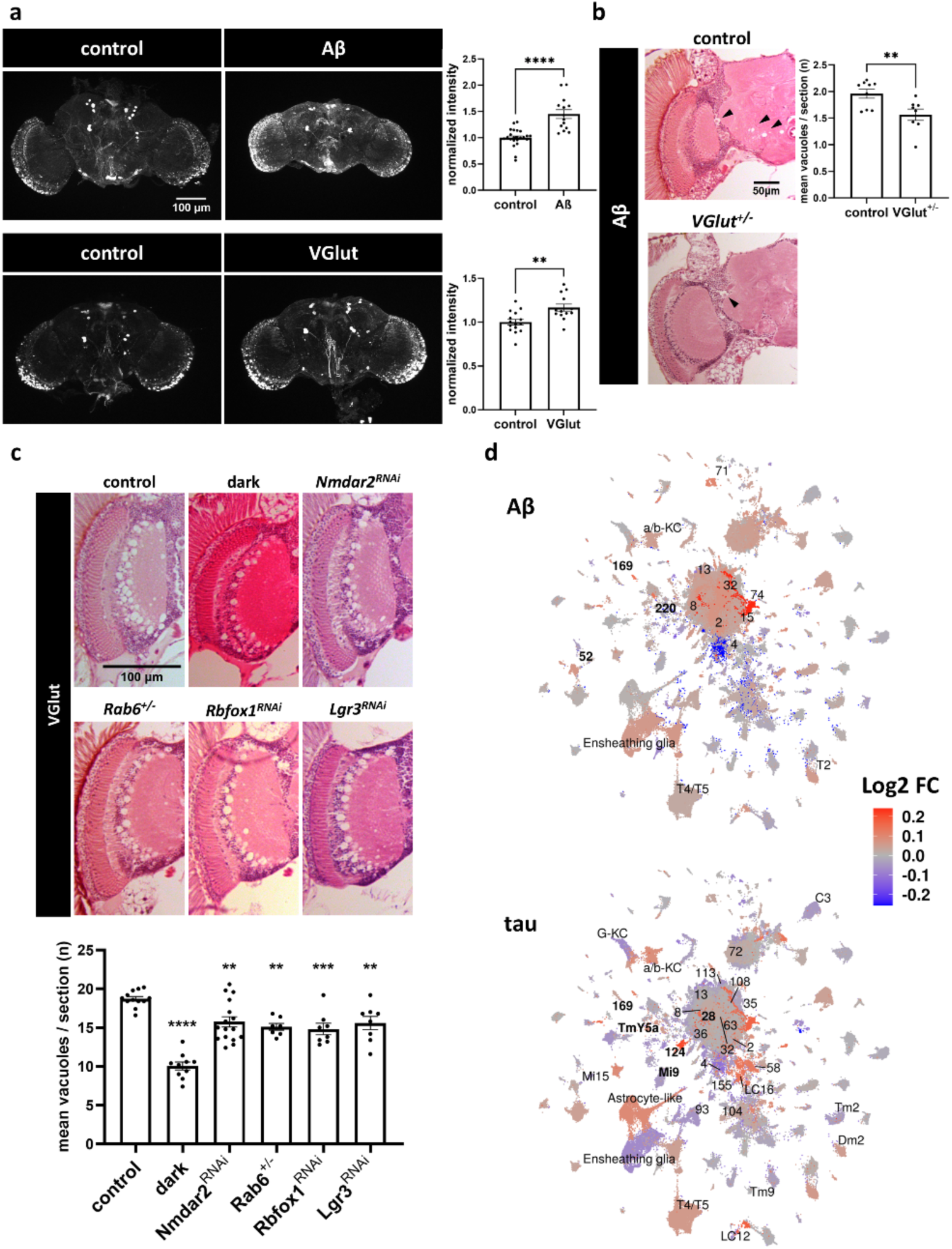
PHGbrown modulates glutamate-induced hyperexcitability. **a**, Elevated Ca^2+^ (anti-GFP, white) is detected in the *Drosophila* brain following pan-neuronal expression of Aβ (*elav-Gal4/+; TRIC-GFP/+; UAS- Aβ42 /+*) versus control (*elav-Gal4 / +; TRIC-GFP / +*), based on whole-mount immunofluorescence of adult brains at day 10 (Z-stack with maximum intensity projections are shown). The VGlut overexpression model (*VGlut-Gal4 / +; UAS-VGlut / TRIC-GFP*) also shows elevated Ca2+ (4-day-old animals), consistent with glutamate-induced hyperexcitability when compared with controls (*VGlut-Gal4 / +; TRIC-GFP / +*). Full genotype of the *TRIC-GFP* reporter system: *TRIC-GFP: LexAop2-mCD8::GFP, nSyb-MKII::nlsLexADBD, QUAS-p65AD::CaM / +; nSyb-QF2 / +* GFP signal intensity was quantified in at least n=13 replicate samples, with statistical analysis based on two sample t-tests. Data points represent individual biological replicate samples. Error bars denote the standard error of the mean. **, p < 0.01; **** p < 0.0001. **b**, Introducing one copy of a *VGlut* loss-of-function allele dominantly suppresses Aβ-induced degeneration in the adult *Drosophila* brain. Vacuoles (arrowheads) were quantified in at least n = 8 animals at 15 days of age. Statistical analysis based on two sample t-test. Each data point represents an individual biological replicate sample. Error bars denote the standard error of the mean. **, p < 0.01. Consistent results were obtained following *VGlut* knockdown using RNA-interference (Extended Data Fig. 10a). **c**, *VGlut* overexpression (*VGlut-GAL4 / +; UAS-VGlut / +*) induces activity-dependent, vacuolar degenerative pathology, and is suppressed by manipulations that reduce activity of PHGbrown driver gene homologs. The following RNAi knockdown or loss-of-function alleles were tested in heterozygosity: *Nmdar2(NIG14794R-3III); Lgr3(v330603)*; *Rab6(08323)*; and *Rbfox1(NIG32062Ra-3)*. Vacuoles were quantified in at least n = 7 animals at 2 days of age. Statistical analysis is based on one-way ANOVA followed by Dunnett’s test for multiple hypothesis correction. Each data point represents an individual biological replicate sample. Error bars denote the standard error of the mean. **, p < 0.01; ***, p < 0.001; **** p < 0.0001. See also Extended Data Fig. 10b. **d**, Differential expression of the *Drosophila* fM1 module, homologous to PHGbrown, varies by cell-type and in response to Aβ versus tau. Uniform Manifold Approximation and Projection (UMAP) plots showing mean fM1 gene differential expression (log_2_ fold change) across distinct brain cell type clusters, based on single nucleus RNA-sequencing in 10-day-old Aβ (*elav-Gal4 / +; UAS-Aβ42 / +*) or tau (*elav-Gal4 / +; UAS-tau / +*) transgenic flies versus controls (*elav-Gal4 / +*). Cell clusters with significant expression changes (Wilcoxon rank sum test p<0.05) are labeled, with glutamatergic neuron clusters noted in bold. See also Extended Data: Fig. 12 and Table 17.

Since Aβ has been implicated to cause myriad deleterious effects in the brain, we next sought a more restricted, direct model of hyperexcitation brain injury. We adopted a previously published strain in which a *UAS-VGlut* transgene is activated within the endogenous *VGlut* expression domain via a *VGlut-GAL4* driver^60^. These flies manifest increased glutamatergic excitatory neurotransmission, owing to excess glutamate packaging into synaptic vesicles, causing elevated neuronal activity, progressive neurodegeneration, and reduced survival^61^. We confirmed hyperexcitation and neurodegeneration in the *VGlut* overexpression model using TRIC imaging and brain histology, respectively (Fig. 6a,c and Extended Data Fig. 10b). In the brains of 2-day-old adult flies, we noted confluent vacuolar pathology in the outer medulla, which participates in visual processing and receives strong glutamatergic input from pathways originating with photoreceptors in the *Drosophila* retina. This degenerative pattern was strongly suppressed when flies were raised in complete darkness, consistent with activity-dependent excitotoxic brain injury (Fig. 6c). We found that loss-of-function manipulations in several PHGbrown driver genes, including *GRIN2B/Nmdar2, RAB6/Rab6, RBFOX1/Rbfox1,* and *RXFP1/Lgr3*, also significantly suppressed vacuolar degeneration following *Vglut-*induced excitotoxicity (Fig. 6c). Our results are consistent with a model in which decreased expression of the PHGbrown module attenuates hyperactivation neuronal injury in AD.

To determine if hyperexcitability can directly mobilize a transcriptional response similar to PHGbrown, independent of AD pathology, we performed RNAseq profiling in the *VGlut* overexpression model at the same 2-day old time point. Our analyses identified 2,864 significantly differentially expressed genes, including enrichment for regulators of synaptic signaling (GO:0099536, p=2.2×10^−14^, Extended Data: Fig. 11 and Tables 15, 16), and this transcriptional signature significantly overlapped with both human PHGbrown (p=7.5×10^−6^) and the homologous *Drosophila* fM1 coexpression network (p=1.9×10^−14^) (Extended Data Fig. 12e). Unexpectedly however, the direction of differential expression was flipped, with *VGlut-*induced hyperexcitability increasing expression of most synaptic genes, including *GRIN2B/Nmdar2* and several other driver gene modifiers from PHGbrown (Extended Data: Fig. 11c,d and Table 16). By contrast, the PHGbrown coexpression module is predominantly down-regulated in AD postmortem bulk brain tissue (Fig. 4 and Extended Data Fig. 8). In our complementary analyses of the *Drosophila* fM1 synaptic coexpression network (above), we similarly found that gene expression was significantly decreased during aging and this was further reduced in *Elav>tau* flies (Fig. 2b and Extended Data Table 8). Although a similar overall trend was documented in *Elav>Aβ* animals (p=0.059, Extended Data Fig. 3), our longitudinal data hints at a potentially more complex, dynamic pattern (Extended Data Fig. 3). Since AD has differential impact across heterogeneous neuronal subtypes, we further reasoned that many transcriptional perturbations might be masked or attenuated in bulk RNAseq data. Therefore, to confirm and define cell-type specificity, we interrogated single nucleus RNAseq data generated from 10-day-old *Elav>Aβ* flies, when elevated intracellular calcium levels also suggest hyperexcitability. Interestingly, Aβ triggered a strong increase in the fM1 coexpression module expression in numerous cell types, including several excitatory glutatmatergic and cholinergic neuron clusters, and similar changes were seen for PHGbrown gene homologs (Fig. 6d and Extended Data Fig. 12a). By contrast, tau predominantly induced down-regulation for these genes among most of the same cell types; although, selected excitatory neuron clusters showed the opposite pattern similar to Aβ. Lastly, we interrogated a recently-published snRNAseq atlas from 427 human postmortem brains, including many of the same brain autopsies from the AMP-AD bulk RNAseq meta-analysis^62^. Similar to *Drosophila*, we find heterogeneous responses across diverse human brain cell types, including significant up-versus down-regulation of PHGbrown genes in many excitatory neuron subtypes in association with AD clinicopathologic traits (Extended Data: Fig. 12c and Table 17). Therefore, while AD pathology and hyperexcitability have overlapping gene expression signatures, our cross-species analyses reveal that these transcriptional programs may be transformed by aging and disease progression in a cell-type specific manner.

## DISCUSSION

We have deployed a cross-species, systems genetics approach to probe the causal chain between AD pathology, global transcriptional changes, and progressive CNS dysfunction. Among 30 consensus AD coexpression modules, we dissect the conserved transcriptional responses to Aβ, tau, alpha-synuclein, and/or aging—together, these triggers account for the majority of AD-associated gene regulatory changes in human brains. Further, through systematic genetic manipulations and behavioral screening, we pinpoint 141 causal driver genes with *Drosophila* homologs that can enhance or suppress Aβ- or tau-mediated neurotoxicity. The STGblue module, implicated in innate immunity, is strongly up-regulated in AD, promotes neurodegeneration based on the results of fly genetic manipulations, and is significantly enriched for AD genetic risk. On the other hand, the PHGbrown module, including regulators of synaptic transmission, is down-regulated in AD postmortem human bulk brain tissue and enriched for loss-of-function suppressors of Aβ/tau. Our findings further suggest that dynamic changes in these synaptic regulatory genes respond to CNS hyperexcitability and modulate downstream AD brain injury. Overall, our strategy highlights how analyses of human data combined with genetic manipulations in more tractable, experimental models, can functionally dissect AD human brain gene expression networks to pinpoint targets that may amplify or protect against disease.

Based on their common transcriptional signatures in humans and *Drosophila* models, aging and AD pathologic triggers appear to have broadly conserved impact on the brain. Whereas the human brain can only be profiled cross-sectionally following death, *Drosophila* models enable interrogation of the transcriptome and behavior longitudinally, enabling dynamic, multi-scale models of brain gene expression and function, in both the presence or absence of Aβ or tau. We have thus generated a unique and powerful longitudinal, cross-species molecular atlas for AD/ADRD gene expression. We have also created a web-based tool making these data broadly accessible for further exploration and analyses (http://flynda.nrihub.org). One major challenge for gene expression analysis in human postmortem brain tissue is the common occurrence of mixed pathologies. Based on prospective, autopsy series, more than half of clinically-diagnosed AD cases are characterized by the presence of alpha-synuclein Lewy bodies, which may modify AD clinical manifestations^63^. In combination, we find that Aβ, tau, and alpha-synuclein together account for up to 62% of AD-associated gene expression changes, with the proportion of changes uniquely responsive to each pathology varying across the AMP-AD modules. Aging was an even stronger driver of brain gene expression changes. Together, aging and AD/ADRD pathologic triggers collectively account for up to 86% of gene expression changes within AMP-AD modules. The residual unexplained gene expression signatures may relate to other pathologies that cause dementia and commonly co-occur with AD, including TDP-43 and cerebrovascular lesions^63^.

Synaptic dysfunction and loss occur early in AD and are strongly linked with initial clinical manifestations and disease progression^64–66^. In addition, a growing body of work suggests that AD is associated with hyperexcitability due to synapse and circuit level pathology, likely contributing to cognitive impairment and risk for seizures^67,68^. Gene expression profiles from human postmortem bulk brain tissue reveal prominent signatures of reduced synaptic and neuronal markers^2,69^. PHGbrown was the top-ranked module based on its strong enrichment for causal modifiers of Aβ and tau-mediated neurotoxicity in *Drosophila*. Importantly, these results suggest that rather than simply a marker of cell death or synaptic loss, transcriptional perturbations affecting regulators of synaptic transmission may modulate AD pathophysiology. Indeed, our findings reveal a causal chain linking AD pathology, hyperexcitability, dynamic gene expression changes, and neurodegeneration. First, we establish that Aβ causes elevated calcium influx, consistent with neuronal hyperexcitability in the adult *Drosophila* brain. Second, we find that either expression of Aβ or more direct induction of glutamate excitotoxicity via *VGlut* overexpression triggers a transcriptional signature that strongly overlaps with PHGbrown. These gene expression changes appear to be dynamic, with either up- or down-regulation depending on the model (Aβ vs. tau), cell type, and/or time course of disease progression. Thus, the prominent down-regulation of PHGbrown in the AMP-AD bulk RNAseq meta-analysis may reflect the dominant neuronal signatures present at later AD stages^2^. Third, experimental manipulations in *Drosophila* that reduce PHGbrown driver genes also suppress excitotoxic degenerative pathology in the brain. We therefore propose a biphasic model in which AD pathology causes an early activation of PHGbrown that promotes neuronal injury followed by a decrease in expression that may be compensatory. Since Aβ aggregation into amyloid plaques precedes the formation of tau neurofibrillary tangles^70,71^, it is possible that the dynamic progression of PHGbrown changes could in part correspond in part to the early Aβ and later tau phases of disease. Complementary analyses of mouse AD models, including APP transgenics, are potentially consistent with an early up-regulation in AMP-AD coexpression modules enriched for neuronal/synaptic markers^2,72^. Additional studies will be required, however, to further elaborate and confirm key details. More granular, longitudinal transcriptome profiles, particularly from AD mouse models, are needed to definitely resolve the time course of gene expression, including both bulk and snRNAseq to reveal both global and cell-type specific changes. The aging brain likely comprises a complex transcriptional mosaic, with signatures from distinct cell types corresponding in part to different disease stages, shaped by the pattern of AD pathologic progression and relative vulnerability versus resilience.

Toxic Aβ oligomers have been shown to directly increase glutamate release^56,73,74^ and/or block reuptake^75^, leading to CNS hyperexcitability in AD. Signaling by NMDA-type glutamate receptors appear to be a central mediator^57^, and our cross-species screening strategy validated *GRIN2B* as a PHGbrown causal driver. Genetic manipulations reducing excitatory neurotransmission, including loss-of-function in either the *Drosophila GRIN2B* homolog, *Nmdar2*, or *VGlut*, which loads glutamate into synaptic vesicles, similarly suppressed Aβ-induced neurodegeneration. Notably, the NMDA antagonist, memantine, which directly binds and targets GRIN2B, is an FDA-approved AD therapy developed to counteract excitotoxic neuronal injury^76,77^. While memantine has demonstrated significant benefit in multiple large clinical trials^78–80^, it provides only modest and transient, overall symptomatic benefit. It is possible that targeting glutamatergic neurotransmission would be more effective at earlier, pre-symptomatic AD stages. Alternatively, memantine efficacy may be transient and limited since *GRIN2B* may already be down-regulated in the brain as part of a compensatory transcriptional response. Regardless, since *GRIN2B* is only one of more than 2,000 genes in the PHGbrown module, more robust neuroprotection may require simultaneous engagement of multiple drivers. In this manner, results from our screen may serve to guide the development of more effective, network-based AD therapeutics.

Our study is notable for several strengths. First, we begin with a robust set of 30 AMP-AD consensus coexpression modules that are each strongly associated with AD and derived from studies of more than 2,000 human postmortem tissue samples^2^. We nominated candidate driver genes based on independent bioinformatic criteria, and our results suggest that these 3 strategies are complementary, yielding additional causal genes in combination. Second, our screen employed established *Drosophila* transgenic models of Aβ, tau, and alpha-synuclein that have been extensively characterized and have proven utility for understanding AD/ADRD^28,29^. Our longitudinal study design, both for *Drosophila* gene expression profiles and genetic screening, embraces the importance of aging as an AD risk and modifying factor. Third, the *Drosophila* locomotor behavioral assay is highly sensitive to determinants of progressive CNS dysfunction that may precede neuronal loss^81,82^, thereby highlighting early causal mechanisms. We also acknowledge some important limitations. In order to accomplish a systematic survey of all 30 AMP-AD coexpression modules, we limited our screen to prioritized, candidate drivers, and it may be informative to probe a selected network in greater depth. In addition, networks based on bulk brain RNAseq may obscure important cell-type specific transcriptional changes, as exemplified for PHGbrown. Given the importance of neuroglial interactions in AD pathophysiology, it will also be important to consider systematic manipulations of candidate drivers in other cell types besides neurons. Nevertheless, our cross-species, systems genetic strategy begins to functionally dissect the complex transcriptional landscape in human brains affected by AD, revealing upstream disease triggers and highlighting promising causal networks for potential therapeutic targeting.

## ACKNOWLEDGEMENTS

We thank Dr. Ying-Wooi Wan for bioinformatic support and feedback on network analysis. The results published here are in part based on data obtained from the AMP-AD Knowledge Portal (https://doi.org/10.7303/syn2580853). ROSMAP study data were provided by the Rush Alzheimer’s Disease Center, Rush University Medical Center, Chicago. We also thank the Bloomington *Drosophila* stock center, the Vienna *Drosophila* RNAi Center, the National Institute of Genetics, Japan, and FlyBase.

## METHODS

### Human Subjects

The Accelerating Medicines Partnership-Alzheimer’s Disease (AMP-AD) consortium meta-analysis of AD differential expression and consensus coexpression networks were previously published^2^. In particular, our study utilized the available summary statistics and lists of all reported genes from coexpression modules. We also leveraged published clinical, pathologic, demographic, and RNAseq data from the Religious Orders Study and Rush Memory and Aging Project (ROSMAP). All ROSMAP participants enrolled without known dementia and agreed to detailed clinical evaluation and brain donation at death^52^ Both studies were approved by an Institutional Review Board of Rush University Medical Center (ROS IRB# L91020181, MAP IRB# L86121802). Both studies were conducted according to the principles expressed in the Declaration of Helsinki. Each participant signed an informed consent, Anatomic Gift Act, and an RADC Repository consent (IRB# L99032481) allowing data and biospecimens to be repurposed. We also employed summary statistics from published analyses of ROSMAP data, including differential expression analyses of bulk^27^ and single-nucleus RNAseq^62^. GWAS summary statistics were also used from published analysis of AD^42,43^, Parkinson’s disease^45^, and height^46^.

### *Drosophila* stocks and husbandry

For transcriptome profiling, we used previously characterized *UAS-tau* (ref. 83), *UAS-Aβ42* (ref. 29), and *UAS-αSyn* (ref. 84). The *UAS-tau* is a chromosome III line (*P{UAS-Tau.wt}7B*) encoding a full-length, wildtype human 2N4R MAPT protein isoform. The *UAS-Aβ42* encodes the 42-amino-acid Aβ42 peptide fused to the Argos secretion peptide (*UAS-Aos:hAβ42*; line M17A on chromosome III). The *UAS-αSyn* stock has a recombinant second chromosome with 2 transgenes codon-optimized for *Drosophila,* encoding full-length human alpha-synuclein. Two published^39,85^ transgenic models of Huntington’s disease expressing human huntingtin (*UAS-HTT^NT^*^231^*^Q^*^128^ and *UAS-HTT^FLQ^*^200^) were also included in the longitudinal RNAseq study, and these data contributed to derivation of *Drosophila* coexpression modules. For pan-neuronal expression, we used the *elav^c^*^155^–*GAL4* (ref. 86) driver strain on the first chromosome, which is available from the Bloomington *Drosophila* Stock Center (BDSC; Bloomington, IN, USA). Experimental animals were heterozygous for the *Elav-GAL4* driver and respective transgenes encoding *Aβ42, tau,* or *alpha-synuclein.* For controls, we included both *Elav-GAL4/+* and *w*^1118^. For locomotor testing and RNAseq, female adult flies were aged to 2, 5, 7, 10, 14, 21, 28, 42, and 57, as detailed in Fig. 1b (not all genotypes were examined at all timepoints due to reduced survival). For genetic modifier screening using locomotor behavior, we used the identical *UAS-tau* and *UAS-Aβ42* strains, which were combined in a stock together with *Elav-GAL4*, a heat-shock-induced lethality mutation on the Y chromosome (*P{hs-hid}*) to increase the efficiency of virgin collection^87^, and a balancer carrying *tub-GAL80* to silence the disease associated protein, permitting maintenance of the stock. Virgin females were collected from this stock and crossed to male flies carrying potential modifier alleles. All modifier strains (RNAi and other alleles) showing genetic interactions are detailed in Extended Data Table 11; these lines were obtained from the BDSC, Vienna Drosophila Resource Center (VDRC), or the Japan National Institute of Genetics (NIG). All modifier gene manipulations were tested in heterozygosity. For secondary tests of genetic interactions using adult brain histology, we used the same *UAS-Aβ42* strain, but as in prior publications^88^ we used an alternate *UAS-Tau^R^*^406^*^W^* strain optimized for modifier studies using the brain histology assay. This chromosome III line encodes a 0N4R mutant form of MAPT associated with familial frontotemporal dementia. For histology experiments, we used a control line (*UAS-empty-VK33*) with an empty UAS site at a chromosome III VK33 docking site from Dr. Hugo Bellen). We obtained the following additional stocks from BDSC or VDRC: *P{UAS-VGlut.D}2* (ref. 89); *P{VGlut-GAL4.D}1*(ref. 60); *Df(2L)VGlut[2]* (ref. 90); and *P{GD834}v2574* (ref. 91). The Transcriptional Reporter of Intracellular Ca2+ (TRIC) (ref. 59) was also obtained from BDSC: *w[*]; P{y[+t7.7] w[+mC]=nSyb-MKII::nlsLexADBDo}attP24, P{w[+mC]=QUAS-p65AD::CaM}2, P{y[+t7.7] w[+mC]=13XLexAop2-mCD8::GFP}attP40/CyO; P{y[+t7.7] w[+mC]=nSyb-QF2.P}attP2, P{w[+mC]=tub-QS.P}heph[9B]*. For the locomotor screen for modifiers of tau, flies were raised at 23℃. Otherwise, flies were raised on standard molasses-based *Drosophila* media at 25℃. Flies were maintained in ambient room lighting unless otherwise noted.

### Analysis of AMP-AD coexpression modules

We leveraged published data from the ROSMAP bulk brain RNAseq differential expression analysis^27^ in order to examine overlaps with each AMP-AD coexpression module (Extended Data Table 1). Differentially expressed genes from analyses considering either Aβ/amyloid or tau/tangle pathologic burden were defined based on |log10 (p-value)| > 1.3, and we considered up- and down-regulated genes separately. For statistical analysis, we performed a hypergeometric overlap test, evaluating the overlap of all genes in each AMP-AD module with the AD pathology-associated, differentially expressed gene sets. For the background, we used the number overall total number of shared genes between the AMP-AD and ROSMAP datasets.

Candidate driver genes were prioritized from the AMP-AD coexpression modules using 3 distinct criteria: i) magnitude of differential expression (extreme), ii) average network degree (hub), and iii) correlation with the module first principal component (PC1, average). Each criterion was used to generate a rank-list from each of the 30 AMP-AD modules. “Extreme” genes were ranked using the true effect (TE) differential expression absolute values from the published AMP-AD RNAseq random effects meta-analysis (data available at syn11914606)^2^. “Hub” gene average degree was computed using the igraph R package^92^ (data available at syn10208070). “Average” genes were ranked based on absolute PC1 correlation (data available at syn11940434). Genes were selected from the top 1% of genes ranked by each criterion from each module, with a maximum list size of 30 ranked genes and a minimum list size of 10 ranked genes for each criterion. A maximum of 90 genes and minimum of 30 genes could therefore be prioritized from each module based on the 3 independent criteria. As detailed in the AMP-AD consortium publication^2^, since consensus coexpression networks were derived independently from multiple brain regions, they can be organized into overlapping clusters. We adjusted our final selections of n=400 gene drivers based on both cluster and module size. Cluster selection sizes were weighted using the formula:

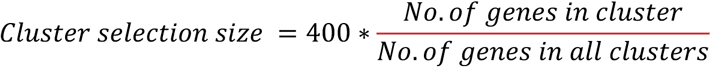

Module selection sizes were then weighted within each cluster using the formula:

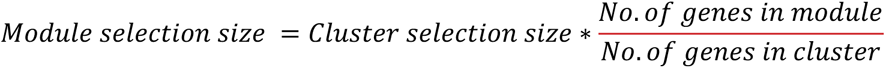

To generate putative causal subnetworks, we first obtained published network adjacency matrices for the AMP-AD modules for each of the 7 brain regions from Synapse (cerebellum/CBE, syn8281722; dorsolateral prefrontal cortex/DLPFC, syn8268669; frontal pole/FP, syn8340017; inferior frontal gyrus/IFG, syn8349785; parahippocampal gyrus/PHG, syn8345109; superior temporal gyrus/STG, syn8343704; temporal cortex/TCX, syn8276546). Each brain region’s adjacency matrix was used to produce network graphs with the *network_from_adjacency_matrix* function in the igraph R package. Prioritized driver gene candidates (or experimentally validated modifier genes) from each AMP-AD module were annotated in their network graphs, and their first-order neighbors were selected using the *neighbors* function in the igraph R package. Subnetworks containing the modifier genes plus their directly coexpressed first-order neighbors were produced for each of the 30 AMP-AD modules using the *induce.subgraph* command, and the resultant undirected subgraph was used for downstream analyses.

The MAGMA tool^44^ was used to conduct gene and gene set enrichment analyses for GWAS signal in the 30 AMP-AD modules and the respective subnetworks, derived using either (i) all prioritized candidate driver genes or (ii) experimentally validated driver gene modifiers (causal subnetworks). GWAS summary statistics were downloaded from studies of AD^42,43^ (NIAGADS: NG00075; https://ctg.cncr.nl/software/summary_statistics), Parkinson’s disease^45^ (https://pdgenetics.org/resources), or height^46^ (https://portals.broadinstitute.org/collaboration/giant/). The *org.Hs.eg.db* R package^93^ was used to map gene symbols to Entrez IDs for each gene in the AMP-AD modules or subnetworks prior to running MAGMA. All analyses with AD GWAS data were repeated without summary statistics from chromosome 19 in order exclude potential confounding due to *APOE*.

Causal mediation analysis examining the relation of PHGbrown and STGblue with AD pathologic burden and cognitive decline were conducted using the *mediation* package in R. These analyses employed clinical, pathologic, and RNAseq (DLPFC bulk brain) data from ROSMAP. For the outcome variable, we use a global cognitive function summary measure (cogn_global). For the predictor variable, we used a summary measure of global AD pathology (globalpath). We computed mean gene expression for all genes in each AMP-AD module as a mediator variable. The age at death (age_death), post-mortem interval (pmi), education (edu) and sex (sex) were included as covariates. For statistical analysis, 1000 simulations were completed using the *mediate()* function.

### Gene expression analysis in *Drosophila* models of aging and AD/ADRD

For longitudinal RNAseq profiles of the *Elav>Aβ42, Elav>tau,* and *Elav>alpha-synuclein* models and controls (*Elav-Gal4* and *w*^1118^), we prepared 3 biological replicates for each genotype and aging time point. For the *VGlut* overexpression model and *VGlut-Gal4* controls, 4 biological replicates were generated. Each replicate consisted of 100 heads collected from flash frozen, age-matched virgin female flies. mRNA was extracted using TRIzol (#15596026, Invitrogen) followed by DNAse treatment. A minimum of 500 nanograms of total DNase-treated extract RNA were used per replicate. Preparation of sequencing libraries, RNA-sequencing, and alignment was performed by the New York Genome Center (longitudinal analysis of *Drosophila* AD/ADRD models) and the Baylor College of Medicine Genomic and RNA Profiling Core (VGlut overexpression model). Briefly, samples were prepared using the Illumina TruSeq Stranded mRNA Library Prep Kit. Samples were then sequenced on an Illumina NovaSeq 6000 with 100 bp paired-end reads. Raw reads were aligned to Drosophila reference genome dm6 r6.06 and quantified. Genes with an average read count < 50 across all samples were excluded. Gene quantification was performed using the featureCounts function from the Rsubread package.

Differential expression analysis was performed using DESeq2 (1.32.0) in R (4.1.0)^94^ (Extended Data Table Table 2). Raw transcript counts were normalized for library depth and composition using the DESeq2 median of ratios normalization approach. To identify gene expression changes during normal aging within control flies (*w1118*), we first modeled age as categorical variable and performed the likelihood-ratio test to determine the significance of the age term (reduced model: expression ∼ 1). To identify gene expression changes triggered by each AD/ADRD pathologic trigger, we employed the following linear model with age as a categorical variable (expression ∼ genotype + age). We tested the significance of the genotype term coefficient using a likelihood-ratio test comparing with a reduced model (expression ∼ age). Functional pathway enrichment analysis was completed using the *G:GOSt* function from the R package gProfiler2 (0.2.1)^95,96^. The Gene Ontology (GO), and KEGG databases were used for querying genes. We used the hypergeometric overlap test (*phyper* function in R) to evaluate the overlap/enrichment of each AMP-AD human coexpresison module with the differential expressed gene sets defined from the tau, Aβ, αSyn models or aging control flies (Fig. 1e,f and Extended Data: Fig. 1c and Table 5). The genes within each AMP-AD human coexpression module were mapped to *Drosophila* using DIOPT^38^ with a DIOPT score filter > 5 (Extended Data Table 4). When more than one fly homolog had a DIOPT score > 5, all were included. For overlap testing, the total number of shared genes between the *Drosophila* RNAseq and AMP-AD datasets (after DIOPT conversion) was used as the background.

### Analysis of *Drosophila* coexpression networks

Weighted gene coexpression network analysis (WGCNA) was implemented as previously described^26^ using the WGCNA package in R^35^. Expression counts from 147 unique *Drosophila* samples (13,192 transcripts) were included in the analysis after normalization in DESeq2 (median-of-ratios depth normalization). The soft power value was set as 4 (scale-free R^2^ = 0.9). To achieve maximum sensitivity, deepSplit was set as 4 and cutHeight was set as 0.1. Minimum module size was set as 23. Closely related modules were merged based on module eigengenes at a distance threshold of MEDissThres = 0.1 (See Extended Data Fig. 2 for the cluster dendrogram). To indicate expression behavior of the modules, module “eigengenes” were calculated with the *moduleEigengenes()* function in the WGCNA package, and we also computed mean module expression by averaging counts of all gene in each module. We examined cell type signatures for each module using markers from a published *Drosophila* brain single cell sequencing dataset^97^ (Extended Data Fig. 2), employing the hypergeometric overlap test (*phyper* function in R. Functional enrichment analysis of Drosophila gene modules was done using the “WebGestaltR” package in R^98^. The enrichMethod parameter was set to “ORA”. The organism parameter was set to “dmelanogaster”. The enrichDatabase was set to ’geneontology_Biological_Process’. In order to determine module conservation and cross-species overlaps, *Drosophila* genes were mapped to their human orthologs using DIOPT^38^ with a filter of DIOPT score > 5. When more than one human homolog had a DIOPT score > 5, all were considered. The hypergeometric test (*phyper*) was next used to compute overlaps between each *Drosophila* coexpression module and each AMP-AD human coexpression module. For the background, we used the total number of shared genes between complete *Drosophila* RNAseq dataset (following DIOPT conversion) and the AMP-AD dataset. Results were plotted using the *pheatmap* package in R^99^.

In order to examine for differential expression of coexpression modules in relation to age or AD/ADRD pathologic triggers, we used linear regression with mean expression of all genes in each module as the outcome variable (Fig. 2b, Extended Data: Table 8 and Figs. 3,4). We considered RNAseq data from each AD/ADRD model (*Elav>Aβ42, Elav>tau,* or *Elav>alpha-synuclein*) along with the *Elav-Gal4* driver control. We compared the full model (module expression ∼ age + genotype) to a reduced model (module expression ∼ age). Age was coded as a categorical, factor trait to account for the possibility of non-linear effects. The genotype term coefficient was used for significance testing using likelihood ratio test. To assess the relationship between module expression and locomotor behavior (Fig. 2c and Extended Data Fig. 5), we first tested the Pearson correlation between module expression and climbing speed using the *cor.test()* function in R. For fM1/fM11, we further implemented linear regression using a base model with locomotor behavior (climbing speed) as the outcome variable, and both age and genotype as predictor variables variables. Mean fM1 or fM11 module expression was then included as an additional predictor variable. To assess for potential mediation, we computed the percentage change in the genotype term β-coefficient (absolute value) with or without inclusion of the term for module expression in the model. The β-coefficient of the genotype term from the full model, including fM1/fM11 expression was considered as the direct effect.

### *Drosophila* genetic modifier screen

For genetic screening, we mapped homology for each prioritized candidate AMP-AD driver using DIOPT^38^. We used a DIOPT score of 5 or greater (out of a maximum possible score of 15), and up to 3 total fly gene homologs were promoted for screening; the top 3 highest scoring genes were selected in cases with more than 3 conserved *Drosophila* homologs. The negative geotaxis climbing assay was performed using a custom robotic system (SRI International, available in the Automated Behavioral Core at the Jan and Dan Duncan Neurological Research Institute) as previously descripbed^39^. The robotic instrumentation elicited negative geotaxis by ‘tapping’ *Drosophila* housed in 96-vial arrays. After three taps, video cameras recorded and tracked the movement of animals at a rate of 30 frames per second for 7.5 s. For each genotype, we collected 4 replicates of 10 females to be tested in parallel (biological replicates). Each trial was repeated five times (technical replicates) per trial day. Flies were transferred into new food every day. The automated, high-throughput system is capable of assaying 16 arrays (1536 total vials) in ∼3.5 hr. To transform video recordings into quantifiable data, individual *Drosophila* were treated as an ellipse, and the software deconvoluted the movement of individuals by calculating the angle and distance that each ellipse moves between frames. Replicates were randomly assigned to positions throughout the plate and were blinded to users throughout the duration of experiments. The results of this analysis were used to compute the average speed of the 10 females in each vial/time point. Software required to run and configure the automation and image/track the videos include Adept desktop, Video Savant, MatLab with Image Processing Toolkit and Statistics Toolkit, RSLogix (Rockwell Automation), and Ultraware (Rockwell Automation). Additional custom-designed software includes Assay Control – SRI graphical user interface for controlling the assay machine; Analysis software bundles: FastPhenoTrack (Vision Processing Software), TrackingServer (Data Management Software), ScoringServer (Behavior Scoring Software), and Trackviewer (Visual Tracking Viewing Software).

The modifier screen used 1,532 alleles targeting 433 *Drosophila* genes that were homologous to 357 human genes. The female progeny from each cross were collected within a 12 hour period. We assessed climbing from 5 to 18 days post-eclosion in our initial locomotor screen. On average 6 timepoints were assessed for the tau model (as it shows an earlier phenotype) and 9 time points for Aβ. As in prior published work^39,100^, we assessed locomotion in *Drosophila* as the average speed at which animals moved in each vial as a function of age and genotype using a nonlinear random mixed effects regression model. ANOVA was used to assess statistical significance followed by Bonferroni-Holm correction to determine adjusted p-values. Specifically, we looked at differences in regression between genotypes, genotypes with time (additive effect, represented by a shift in the curve) or the interaction of genotype and time (interactive effect, represented by a change in the slope of the curve). We estimated the expected statistical power to detect differences by each of our models using a threshold for statistical significance (alpha = 0.05). All plots were reviewed to confirm that statistically significant results from the nonlinear mixed effects models represented meaningful differences on visual inspection. Statistical analysis and graphical visualizations where generated in R^101^. For those strains that enhanced tau- or Aβ-induced locomotor behavior, we performed an additional test to evaluate the consequences of gene manipulation independent of tau/Aβ, crossing with the *elav>GAL4* driver line. Female progeny were then tested for 35 days to determine whether the modifier can be classified as additive or synergistic, based on its toxicity profile in the absence of tau/Aβ (Extended Data Table 11). All modifiers from PHGbrown were retested and locomotor behavior was assayed longitudinally from 3 to 28 days (Fig. 3d and Extended Data Fig. 6).

In order to consider a prioritized candidate driver as an experimentally validated modifier gene, we required consistent results from at least 2 independent allele strains (either consistent, significant enhancement or suppression for a given direction of manipulation, either loss- or gain-of-function). For all such validated modifier genes, we attempted to generate “causal predictions” (amplifying vs. compensatory) by integrating the results from the *Drosophila* locomotor tests (either loss- or gain-of-function enhancer vs. suppressor) with the human differential gene expression (up-versus down-regulated in AD). First, for each modifier gene, we first determined a consensus for the observed interactions based on the results for alleles tested and considering tau versus Aβ separately. Second, we used the consensus results for loss- or gain-of-function of the fly gene to infer the potential impact of differential expression in human brains. For example, if a given human gene is down-regulated in brains affected by AD, and loss-of-function of the fly homolog led to suppression of Aβ/tau neurotoxcity, the gene would be annotated as “compensatory”; whereas, if loss-of-function led to enhancement of Aβ/tau neurotoxcity, the gene would be annotated as “amplifying”. For genes in which the consensus result had opposite interactions in the tau versus Aβ models, we did not assigning any causal predictions (gray nodes in the subnetwork plots; Fig 4a and Extended Data Fig. 7).

### *Drosophila* brain histology

*Drosophila* heads were fixed using 8% glutaraldehyde (Electron Microscopy Sciences) at 4°C for 7-10 days, followed by paraffin embedding and microtome sectioning, as previously described^29^. Serial 5 μm-thick frontal sections (Leica microtome) were prepared from *Drosophila* heads and mounted on slides, followed by hematoxylin and eosin staining to examine brain morphology. We quantified vacuoles greater than 5 μm in diameter in the central brain in all frontal brain sections (∼20 per animal) and computed the average vacuole count per section. All light microscopic images were acquired using a Leica DM 6000 B system. Brain vacuole and nucleus counts for individual *Drosophila* were then used to conduct two-sample unpaired t-tests between genotypes for significance using GraphPad Prism software. For multiple comparisons, we performed one-way ANOVA with Dunnett’s post-hoc test.

### *Drosophila* Calcium Imaging

Immunofluorescence imaging was performed to quantify the TRIC signal. Initiation of TRIC expression via the QUAS system was performed as previously described^102^. Flies carrying TRIC elements were placed in food vials containing 300ul of 250mg/ml quinic acid solution 3 days before they reach the target age. The immunostaining procedure was adapted from published protocols^103^. Flies were anesthetized with CO2 and brains were dissected with forceps in PBS solution (GeneDEPOT, P2101-050) and fixed in 3.7% formaldehyde (J.T.Baker, 2106-01) for 20 minutes in room temperature. Formaldehyde was removed and brains were washed using PBS with 0.3% Triton-X (Sigma-Aldrich, T8787) (PBST) for 15 minutes by 3 times. Primary antibodies were diluted in 0.3% PBST and samples were incubated in primary antibody at 4°C, rocking for 2 days. Primary antibodies were removed, and brains were washed 3 times using 0.3% PBST for 15 minutes each. Secondary antibodies were diluted in 0.3% PBST, and samples were incubated with secondary antibody at 4°C, rocking for 2 days. Secondary antibodies were removed, and brains were again washed 3 times using 0.3% PBST for 15 minutes each. Whole brains were mounted in Vectashield antifade mounting medium (Vector Laboratories, H-1000-10) and stored in the dark at 4°C prior to imaging. Samples were imaged on a Leica Microsystems SP8X confocal microscope. Z-stacks of 1 micron covered the entirety of whole-mount brains. We used the following antibodies and dilutions: Rabbit anti GFP (Invitrogen A11122, 1:500), Mouse anti bruchpilot (DSHB nc18, 1:100), Alexa Fluor® 488 AffiniPure™ Donkey Anti-Rabbit IgG (H+L) (Jackson ImmunoResearch 711-545-152, 1:500), Cy™3 AffiniPure™ Goat Anti-Mouse IgG (H+L) (Jackson ImmunoResearch 115-165-003, 1:500). Quantification of calcium signal was performed using ImageJ software on z-stacks of imaged sections. The entirety of brain region (ROI) of the z-stacked image was selected and regions not containing brain tissue were selected as background. Mean pixel intensity were measured and intensity from the background regions was subtracted from that of ROI for each fluorescence channel. Two sample t-tests were conducted between genotypes for significance using the GraphPad Prism software.

### Single cell RNA-sequencing datasets

A comprehensive description and analysis of the *Drosophila* single cell RNAseq dataset will be published elsewhere. The overall data generation, quality-control, and analysis workflow was adapted from our prior publication^48^ and is also available with the full dataset on synapse (syn34767207). Three biological replicates per genotype were generated, and all animals were aged to 10 days post-eclosion (21 samples total). Each replicate consisted of 16-18 intact, dissected brains from female flies, and were enzymatically dissociated into a single cell suspension as described in Davie et al., 2018. Single cell libraries were prepared per manufacturer’s protocol for the Chromium Single Cell Gene Expression 3’ v3.1 kit (10x Genomics) by the BCM Single Cell Genomics Core. Completed libraries were sequenced by the Baylor Genomic and RNA Profiling Core using the Illumina NovaSeq 6000 platform with a minimum depth of 300,000,000 reads per sample. Illumina BCL files were demultiplexed into FASTQ files by calling the Cell Ranger 4.0.0 mkfastq function. FASTQ files were aligned to the Drosophila reference genome (BDGP6.22.98) and gene counts were quantified using the Cell Ranger 4.0.0 count pipeline. Given the 10x recovery rate estimations, the cell calling algorithm in Cell Ranger was applied by setting the -- expect-cells parameter in count to 10,000 per library.

We extracted the mean expression for all gene members from fM1 or PHGbrown (mapped fly orthologs) for each cell. Cluster-wise Log2 fold-change for the gene set was evaluated comparing *Elav>* model with driver control was calculated by averaging module expression of all cells in the cluster. Significance of expression change was determined using Wilcoxon singed-rank test. Cluster-wise log2 fold change was used to generate UMAP in Figure 6E using the FeaturePlot() function in the Seurat package in R. For the cross-species analysis, we used published human postmortem brain single-cell RNAseq data^62^. Differential expressed genes from analyses of amyloid or neuritic plaque burden (plaq_n) for each cell cluster were tested for overlap with PHGbrown module genes and its subnetworks. The hypergeometric overlap test (*phyper* in R) was applied to examine overlaps between the PHGbrown module gene set and differentially expressed genes among excitatory neuron cell-types, considering either all genes or up-vs. down-regulated genes separately.

### Statistical Analysis

For clarity, all statistical tests are described above alongside the corresponding methods. We also include sample sizes and relevant statistical tests in all figure legends. All statistical analysis relied on two-tailed tests. For all analyses of RNAseq data, p-values were adjusted for multiple hypothesis testing using the Benjamini-Hochberg false discovery rate (FDR), with significance level set at (FDR p-value < 0.05).

### Data Availability

Further information and requests for resources should be directed to and will be fulfilled by Joshua M. Shulman. The previously published data from the Accelerating Medicines Partnership-Alzheimer’s disease (AMP-AD) Consortium are available from the AD Knowledge Portal hosted by Synapse (https://adknowledgeportal.synapse.org), including the consensus coexpression module definitions (https://doi.org/10.7303/syn11932957.1) and AD case/control differential gene expression meta-analysis results (https://doi.org/10.7303/syn11914606). ROSMAP data resources are available from https://www.radc.rush.edu and www.synapse.org. All *Drosophila* bulk brain and single-cell RNAseq data has been uploaded for distribution via synapse (syn34767207). The Fly Longitudinal NeuroDegenerative Atlas (FLYNDA) [http://flynda.nrihub.org] is a web-based application designed to facilitate the analysis and visualization of longitudinal transcriptomic and proteomic datasets from transgenic *Drosophila* models of neurodegenerative diseases, including Alzheimer’s, Parkinson’s, and Huntington’s diseases. The platform allows researchers to explore differential gene expression, pinpoint fly orthologs of human genes, and compare expression levels across multiple disease models, providing valuable insights into disease-associated perturbations and molecular mechanisms. FLYNDA’s user-friendly interface supports customizable visualizations and integrates tools such as MARRVEL and FlyBase for comprehensive cross-species comparisons. The FLYNDA tool will be comprehensively described in a companion publication but is immediately available to facilitate exploration of the data generated as part of this work.

