## Extended Data Figure for "Systems genetic dissection of Alzheimer’s disease brain gene expression networks"

**Fig. 1: Additional cross-species gene expression analysis in *Drosophila* models.**

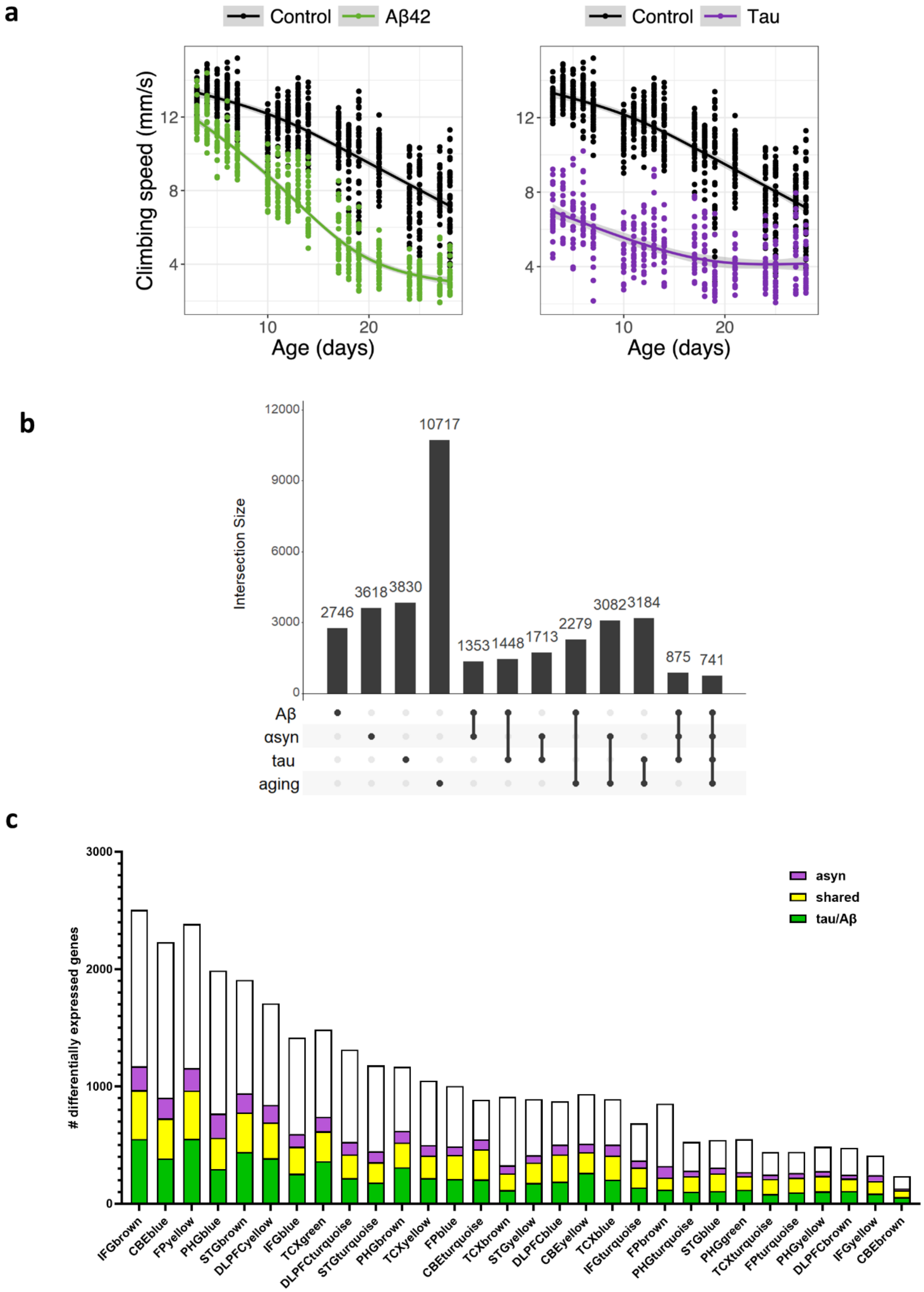


**a**, Pan-neuronal expression of Aβ (*elav-Gal4/+; UAS-Aβ/+*) or tau (*elav-Gal4/+; UAS-tau/+*) cause progressive locomotor impairment when compared with control flies (*elav-GAL4/+*). Statistical analysis based on longitudinal mixed effects regression models and two-way ANOVA [$p_{A\beta} = 4.60 \times 10^{-39}$; $p_{tau} = 3.56 \times 10^{-30}$). Performance was assessed on 18 individual days between day 3 to day 30 of the fly lifespan. Each plotted point represents average speed among a test sample of n=10 flies. Natural spline curve with 3 degrees of freedom was fit to data, and gray shaded area represents the 95% confidence interval for the standard error of the mean. **b**, Upset plot showing number of unique and shared/overlapping differentially expressed genes induced by Aβ, tau, α-synuclein, and aging. **c**, Cross-species analysis highlights genes within human AD-associated coexpression modules, for which expression of conserved *Drosophila* homologs are triggered by Aβ/tau (green), alpha-synuclein (asyn, purple), or both (yellow). Each bar indicates the total module size, based on the total number of conserved genes. See also Fig. 1 and Extended Data Table 5.

**Fig. 2: Weighted gene coexpression network analysis (WGCNA).**

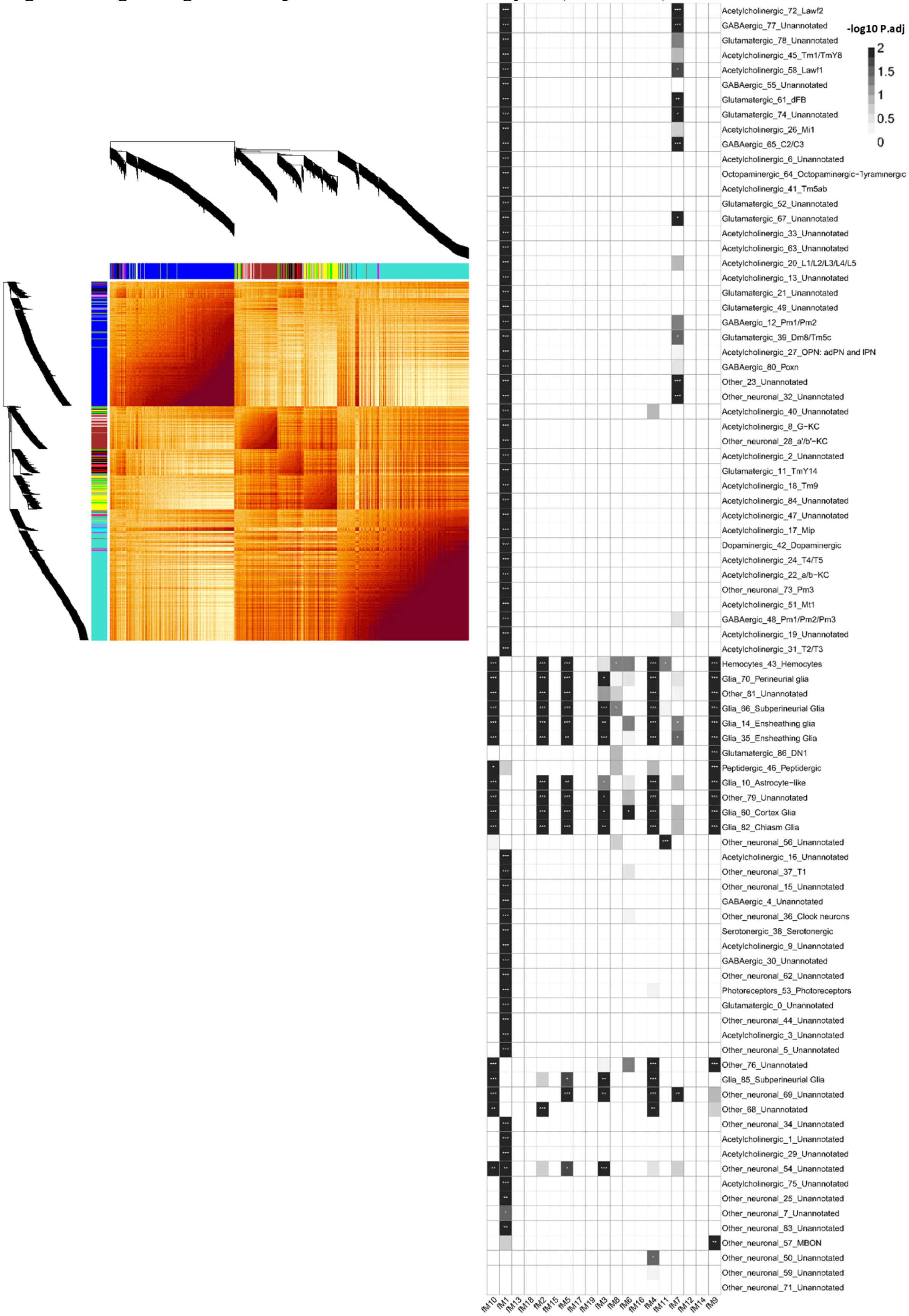


WGCNA heatmap plot (left), with branches in the hierarchical clustering dendrograms and corresponding color bar denoting partitioned modules. Heatmap intensity indicates the strength of co-expression between gene pairs. Heatmap (right) highlighting enrichment of the *Drosophila* coexpression modules (columns) for cell type specific expression signatures (rows), based on the Davie et al, 2018 reference dataset. Gray-scale intensity denotes the significance [$-\log_{10}$ (p-value)] based on the hypergeometic overlap following multiple-hypothesis correction. *, $p < 0.05$; **, $p < 0.01$; *** $p < 0.001$. See also Extended Data Table 2.

**Fig. 3: Dynamic coexpression module changes in *Drosophila* Aβ transgenic models.**

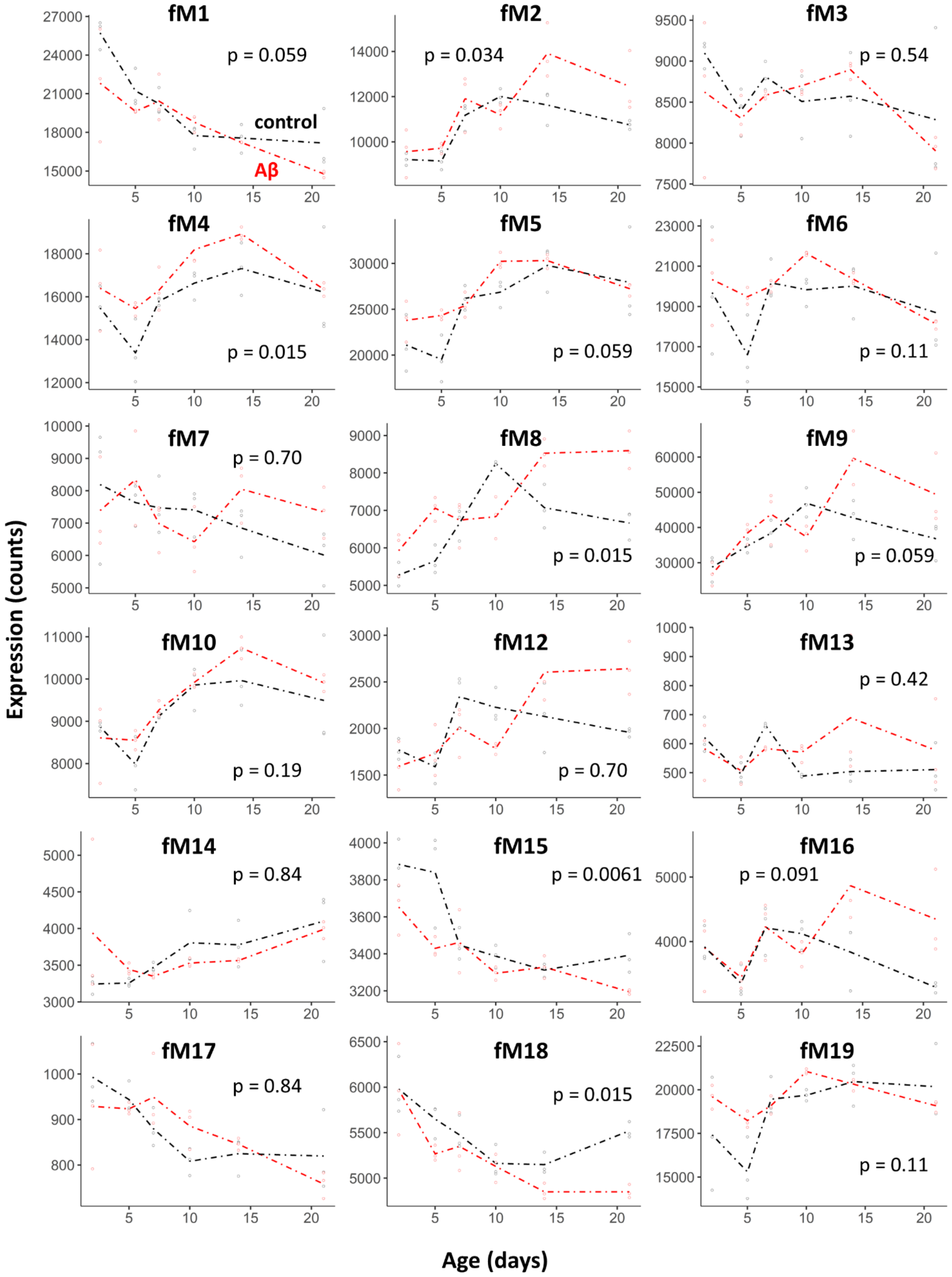


Mean expression for module genes is plotted for the Aβ (red: *elav-Gal4/+; UAS-Aβ/+*) *Drosophila* model and compared with that from controls (black: *elav-GAL4/+*), using linear regression models including longitudinal data (days 2-28 days) and adjusting for age. Statistical analysis based on the likelihood ratio test with significance-level adjusted based on the false discovery rate (*, p<0.05, **, p<0.01). See also Fig. 2 and Extended Data Table 8.

**Fig. 4: Dynamic coexpression module changes in *Drosophila* tau transgenic models.**

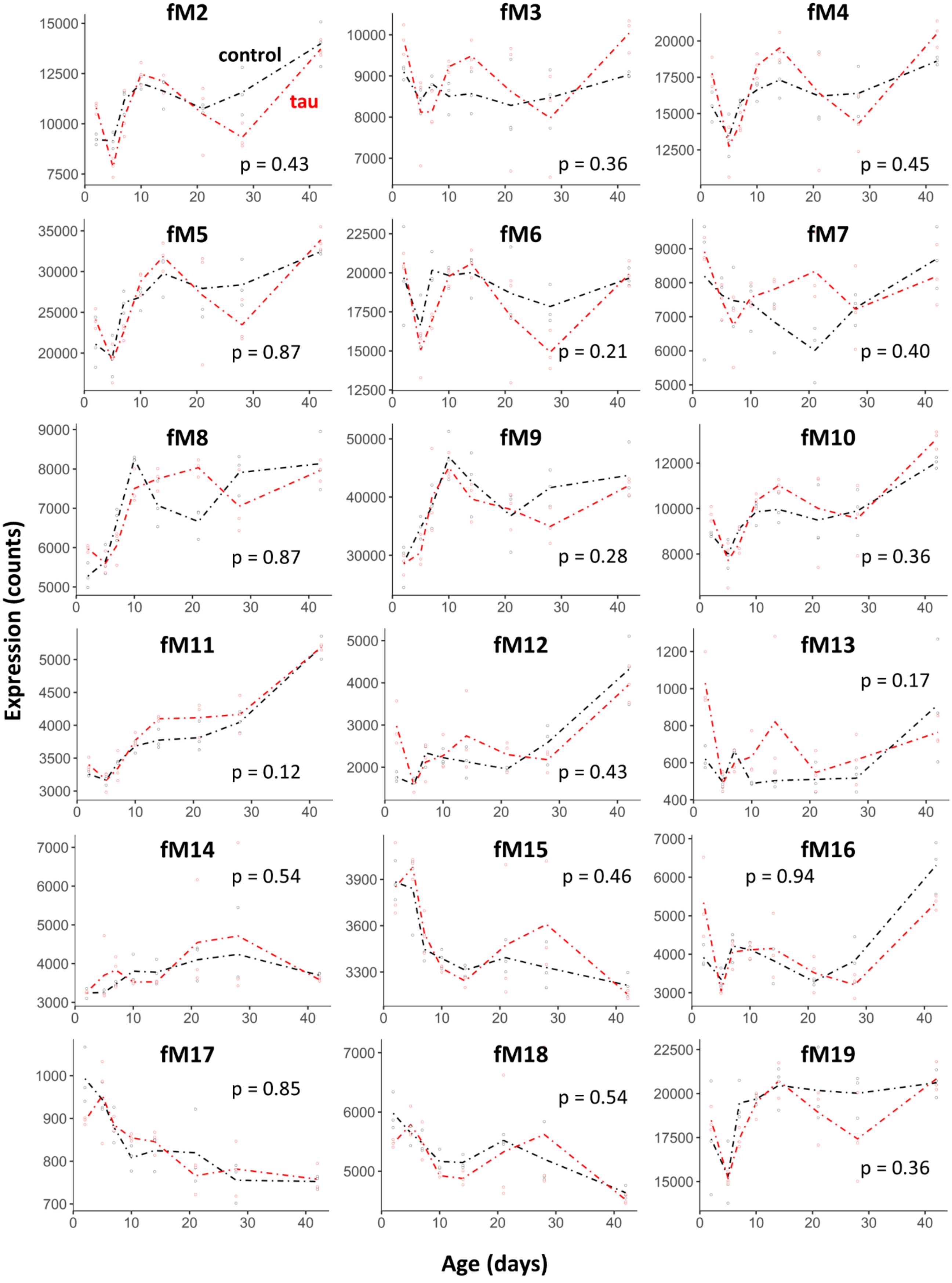


Mean expression for module genes is plotted for the tau (red: *elav-Gal4/+; UAS-tau/+*) *Drosophila* model and compared with that from controls (black: *elav-GAL4/+*), using linear regression models including longitudinal data (days 2-42 days) and adjusting for age. Statistical analysis based on the likelihood ratio test with significance-level adjusted based on the false discovery rate (*, p<0.05, **, p<0.01). See also Fig. 2 and Extended Data Table 8.

**Fig. 5: Integrated analysis of coexpression module expression versus central nervous system function.**

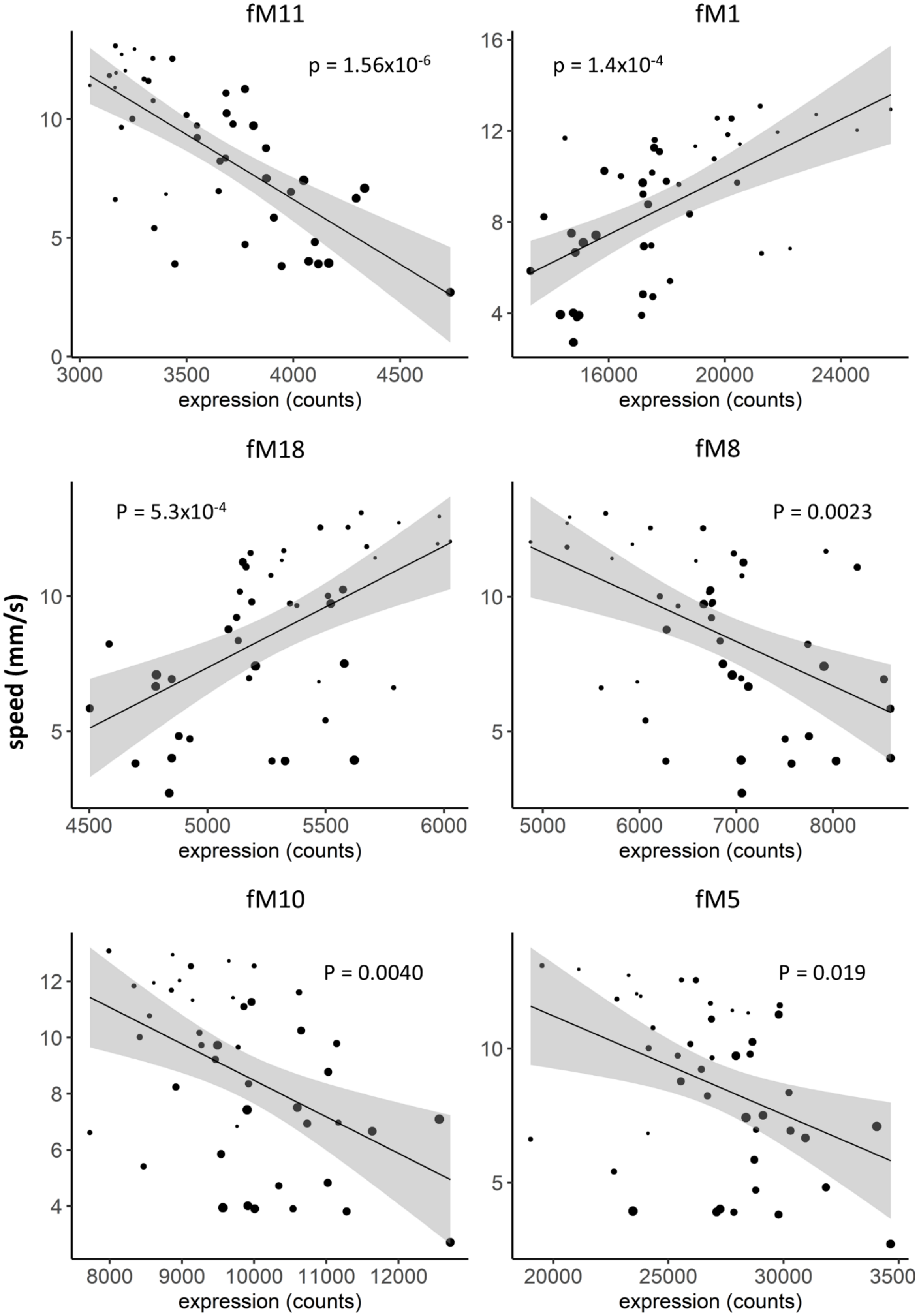


Scatter plots showing relation between mean module gene expression and locomotor behavior (climbing speed), based on Pearson correlation analysis (significance testing based on false discovery rate). The plots include data from 44 different experimental conditions, including 7 genotypes and up to 9 aging timepoints (age denoted by size of the data point). Gray shading denotes the 95% confidence interval (standard error of the mean) for the line of best fit. See also Fig. 2.

**Fig. 6: Additional locomotor screening results for PHGbrown prioritized driver genes.**

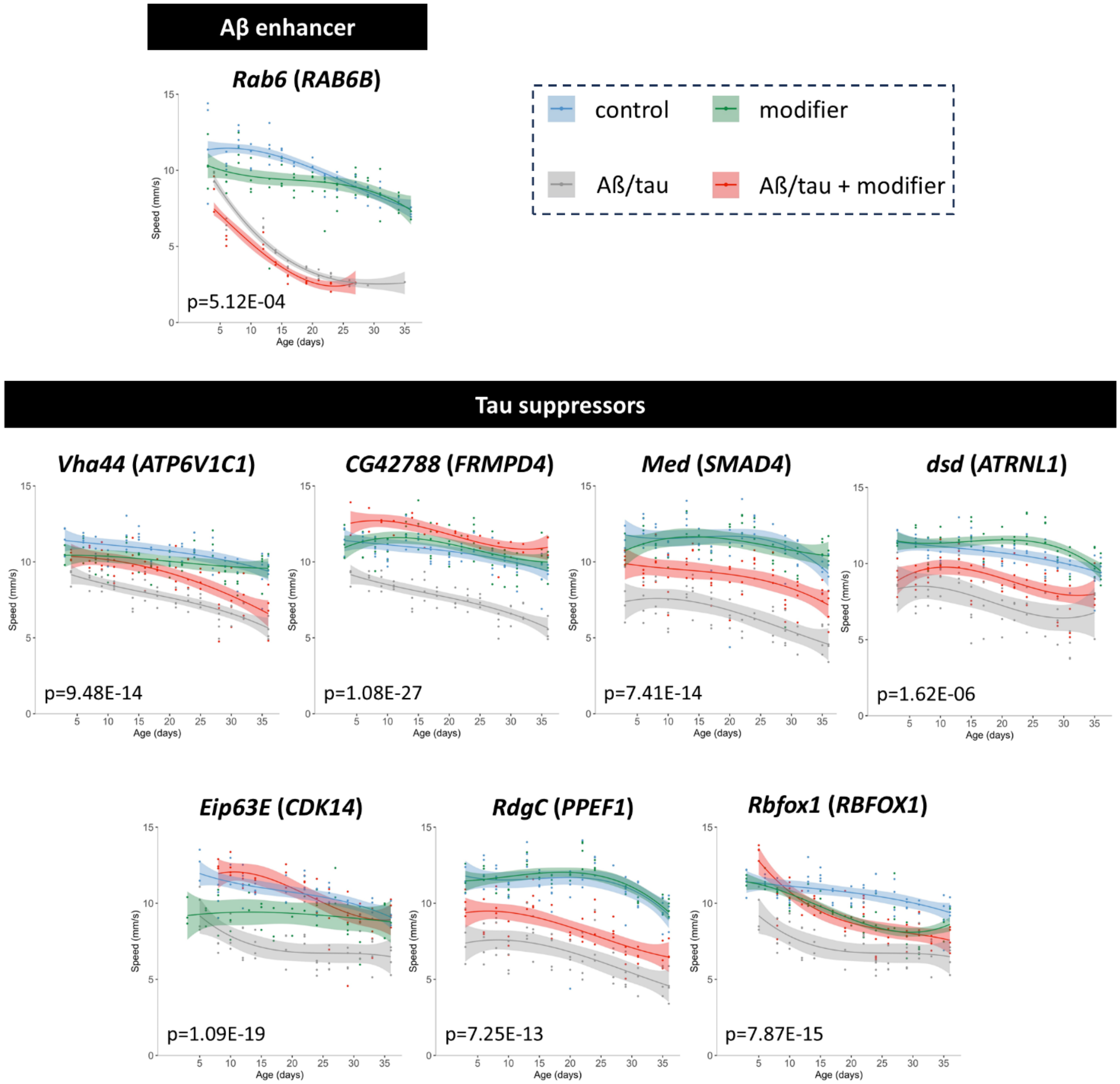


Representative results from tests of additional prioritized PHGbrown candidate driver genes in the Aβ (*elav-Gal4/+; UAS-Aβ42/+*) and tau (*elav-Gal4/+; UAS-tau/+*) transgenic fly models, compared with controls (*elav-Gal4/+*). Plots show the results of longitudinal locomotor behavior assays (climbing speed during startle-induced negative geotaxis). Each modifier test included at least n > 4 replicates of 10 animals each examined at 6 or more timepoints over 35 days of aging. Statistical comparisons based on one-way ANOVA considering three nested models (genotype, genotype + time, and genotype*time) and applying a significance threshold of $p<0.05$, following Holm-Bonferroni adjustment. The following RNAi knockdown or loss-of-function alleles were tested in heterozygosity: *Rab6(GE1331); Vha44(MI02871); CG42788(v45034); Med(v19688); dsd(v1106); Eip63E(MI00413-GFSTF.1); rdgC(v35105);* and *Rbfox1(NIG32062Ra-3II)*. Natural spline curve with 3 degrees of freedom was fit to data, with shading to denote the 95% confidence band. See also Fig. 3 and Extended Data Table 11.

**Fig. 7: Additional Alzheimer’s disease causal gene expression networks.**

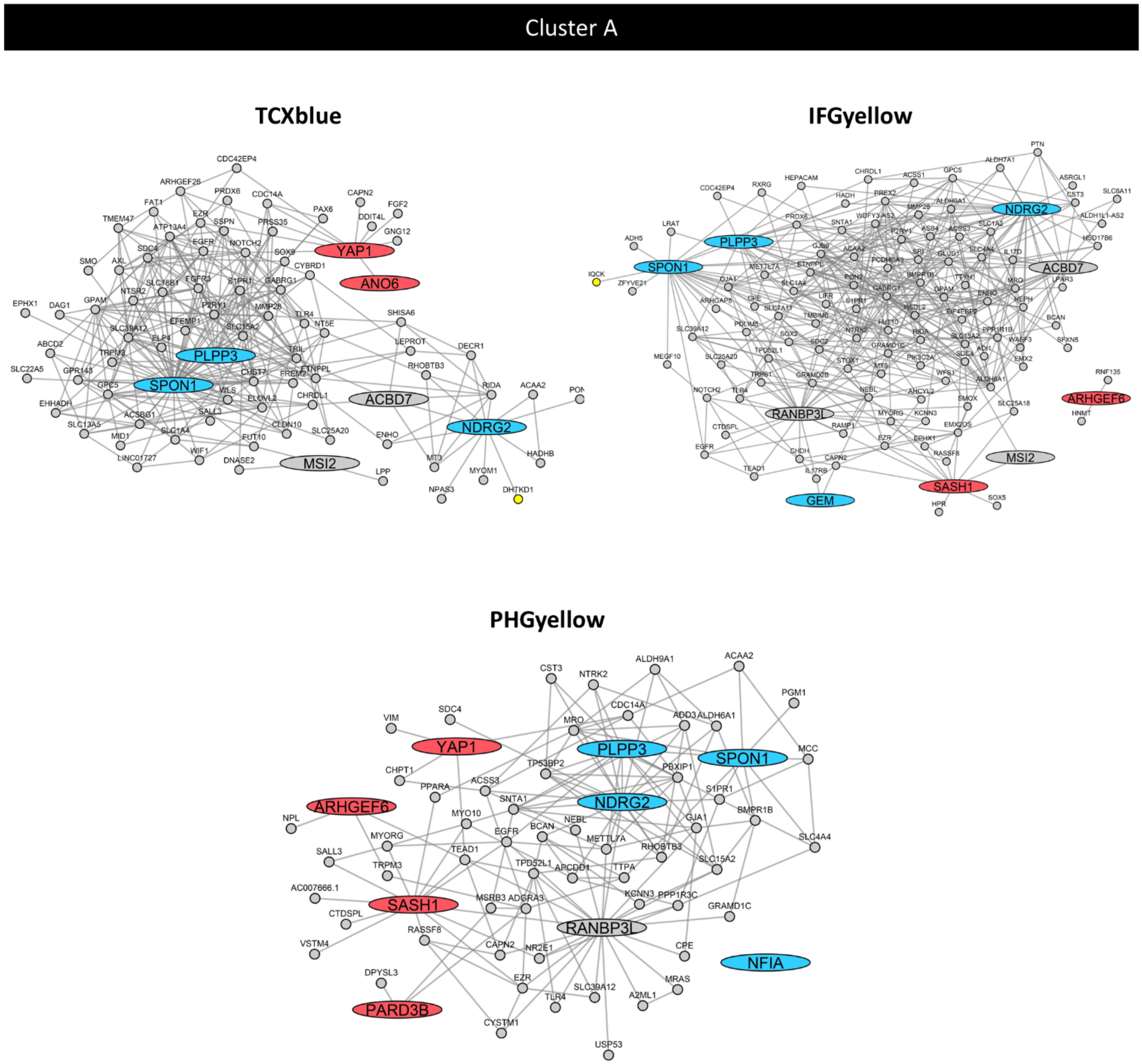


In order to derive subnetworks graphs, we plotted the human homologs of network modifier genes (large oval nodes) along with all first-order neighbors (small circle nodes). Edges indicate coexpression relationships between genes. Up- or down-regulated gene expression changes were based on the published AMP-AD case/control brain transcriptome meta-analysis. Integrating results from the *Drosophila* modifier screen (enhancers or suppressors of Aβ/tau-induced locomotor impairment), we inferred whether causal drivers are predicted to be amplifying or compensatory (red or blue, respectively). Driver genes in gray had data supportive of bi-directional impact. Genes that localize to established AD risk loci are highlighted as yellow nodes. Amplifying gene expression perturbations are expected to promote disease pathophysiology, whereas compensatory changes are predicted to be protective. Subnetworks are organized into 5 overlapping consensus clusters A-E, which are enriched for cell-type specific expression signatures, as follows: astrocytes and pericytes (clusters A and B), endothelial cells (B), microglia (B), neurons (C and E), oligodendroglia (D), and oligodendroglial precursors (A and D). See also Fig. 4 and Extended Data Tables 12-13.

**Fig. 7 (continued)**

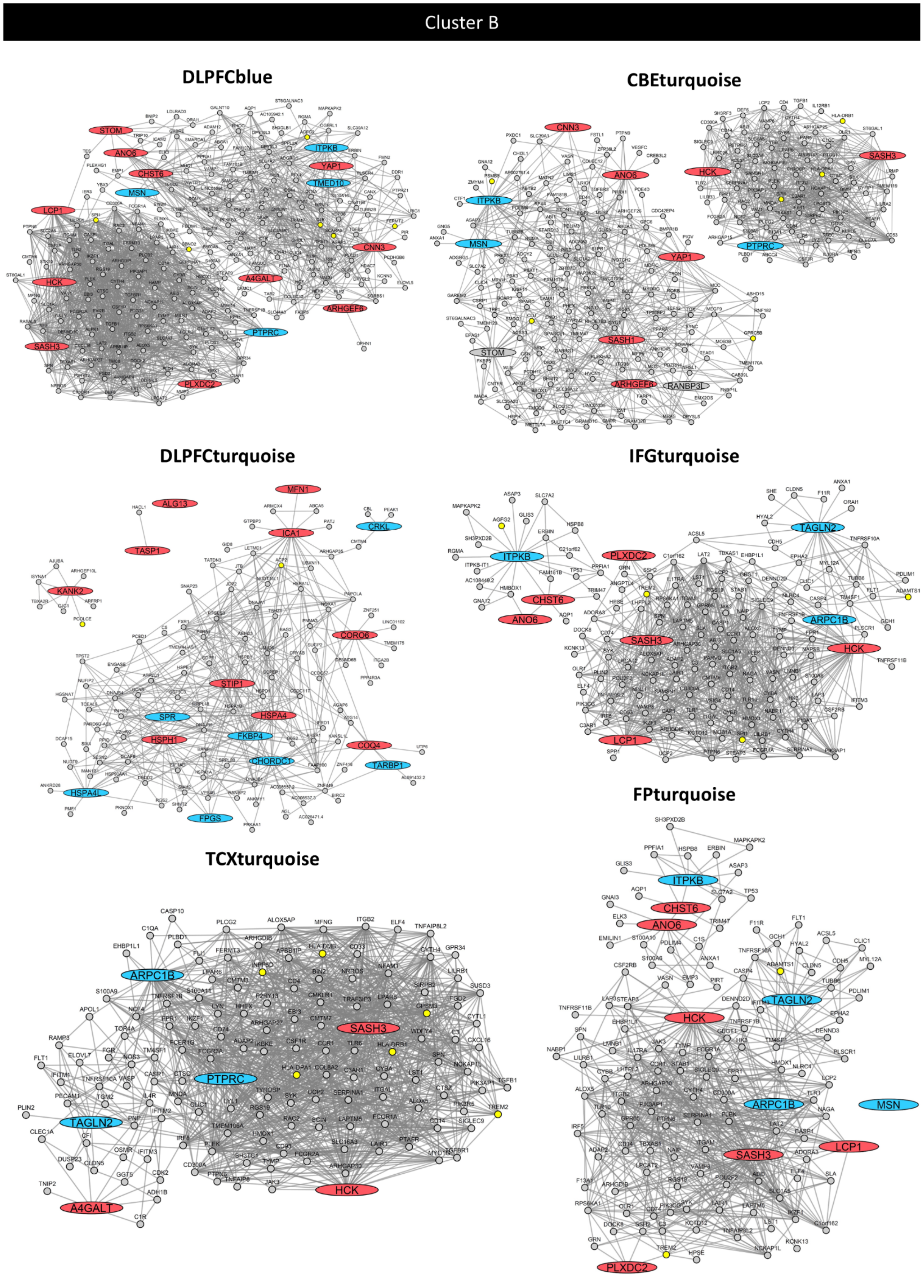

**Fig. 7 (continued)**

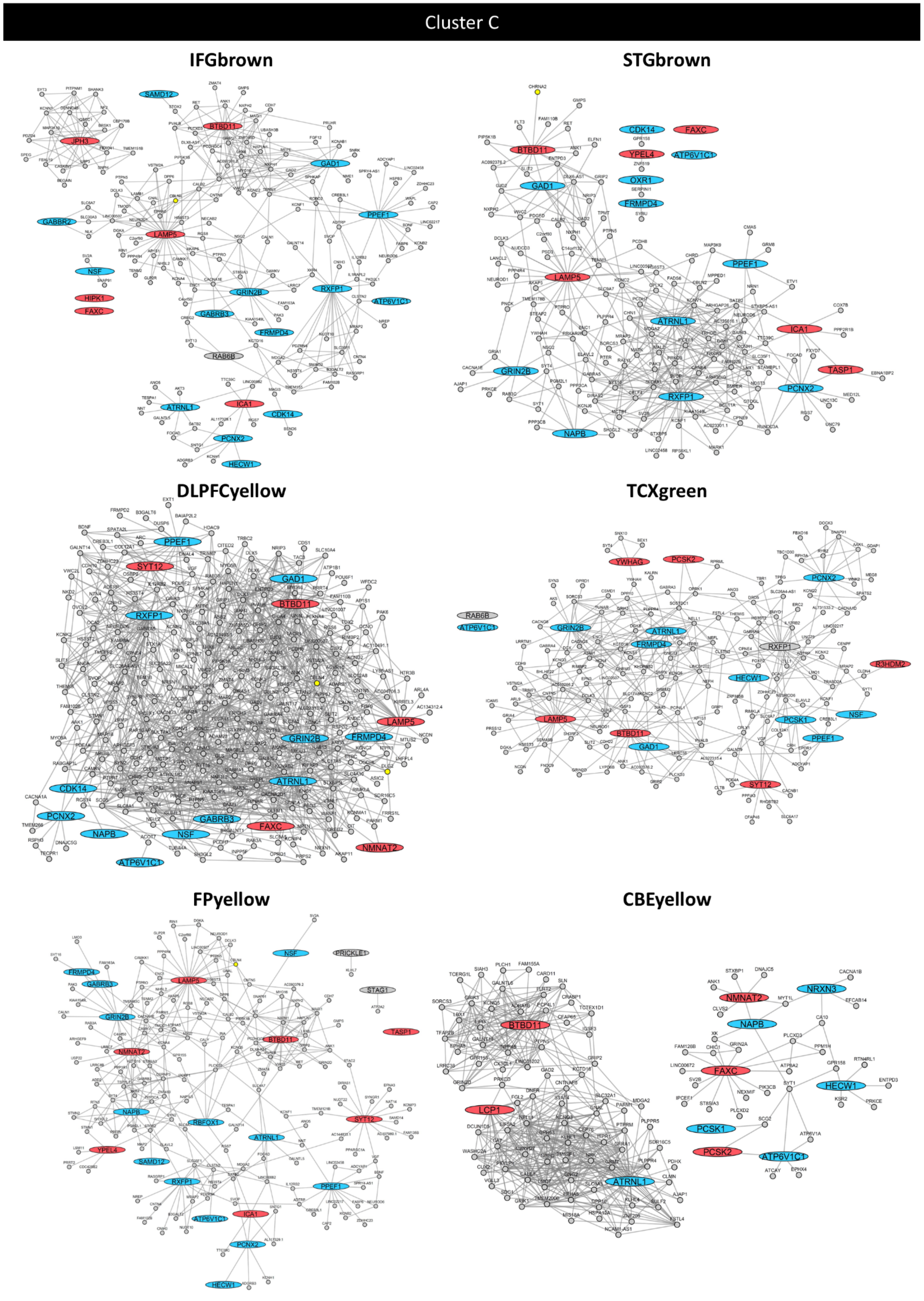

**Fig. 7 (continued)**

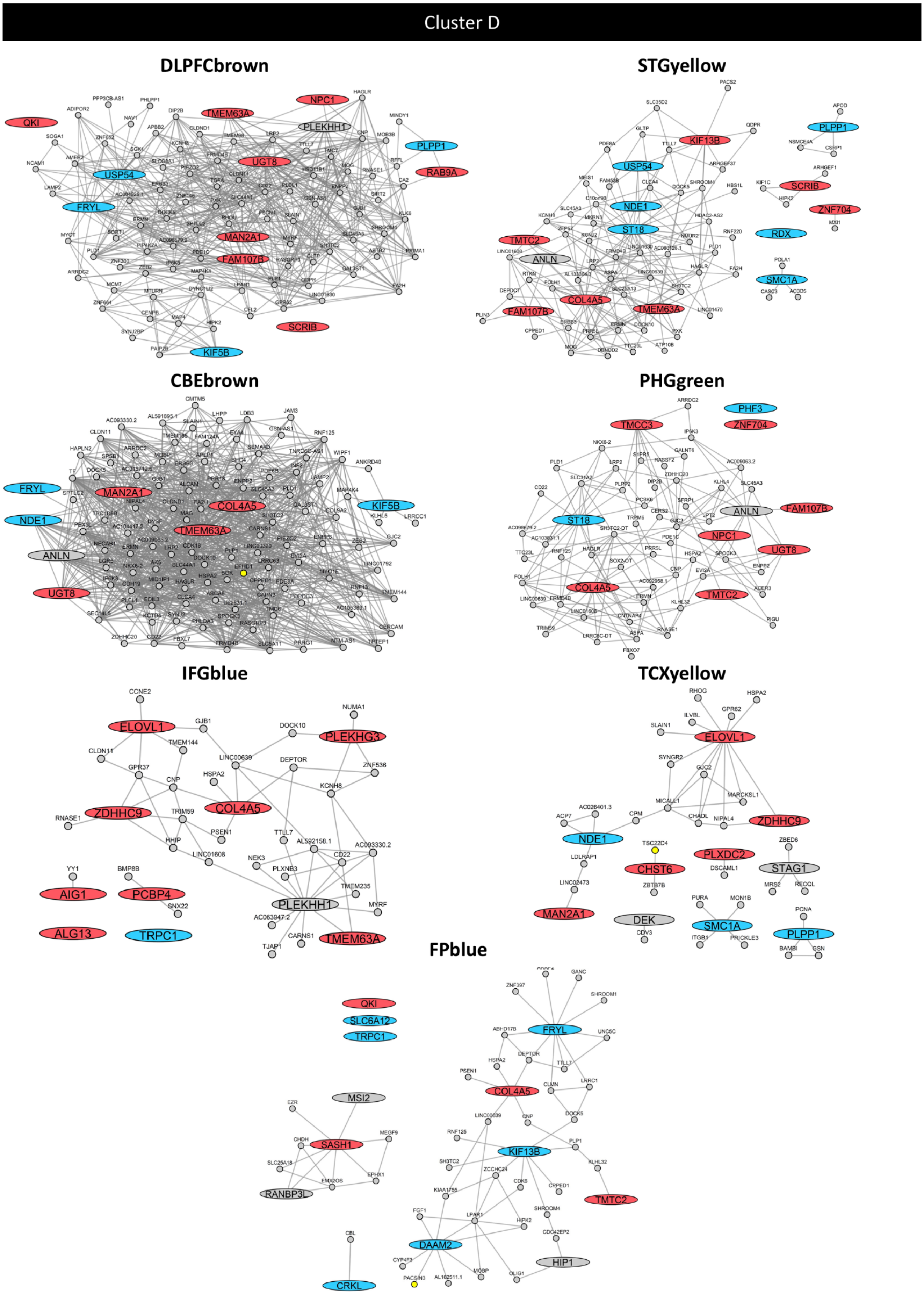

**Fig. 7 (continued)**

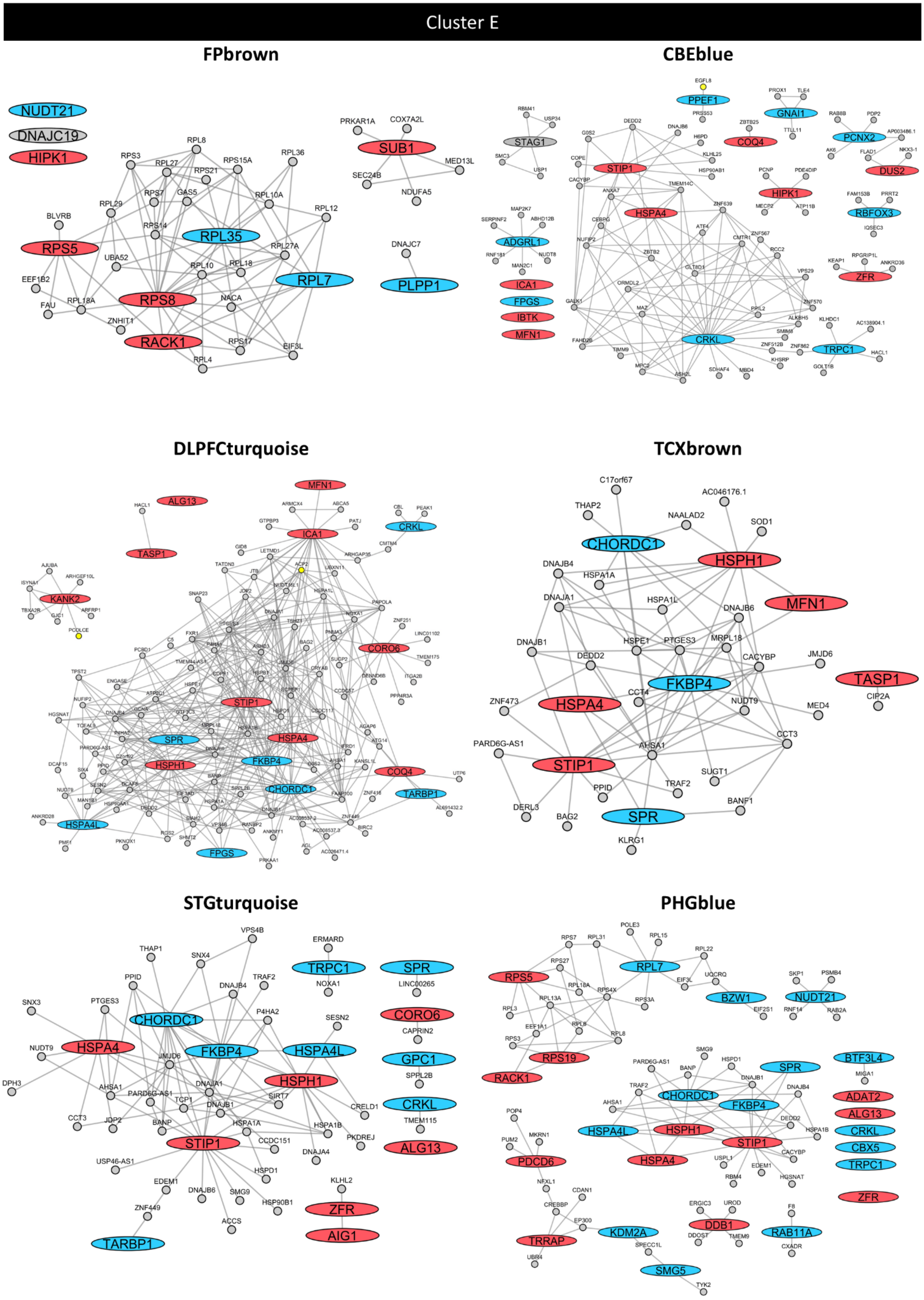

**Fig. 8: The PHGbrown subnetwork is predominantly down-regulated in Alzheimer's disease.**

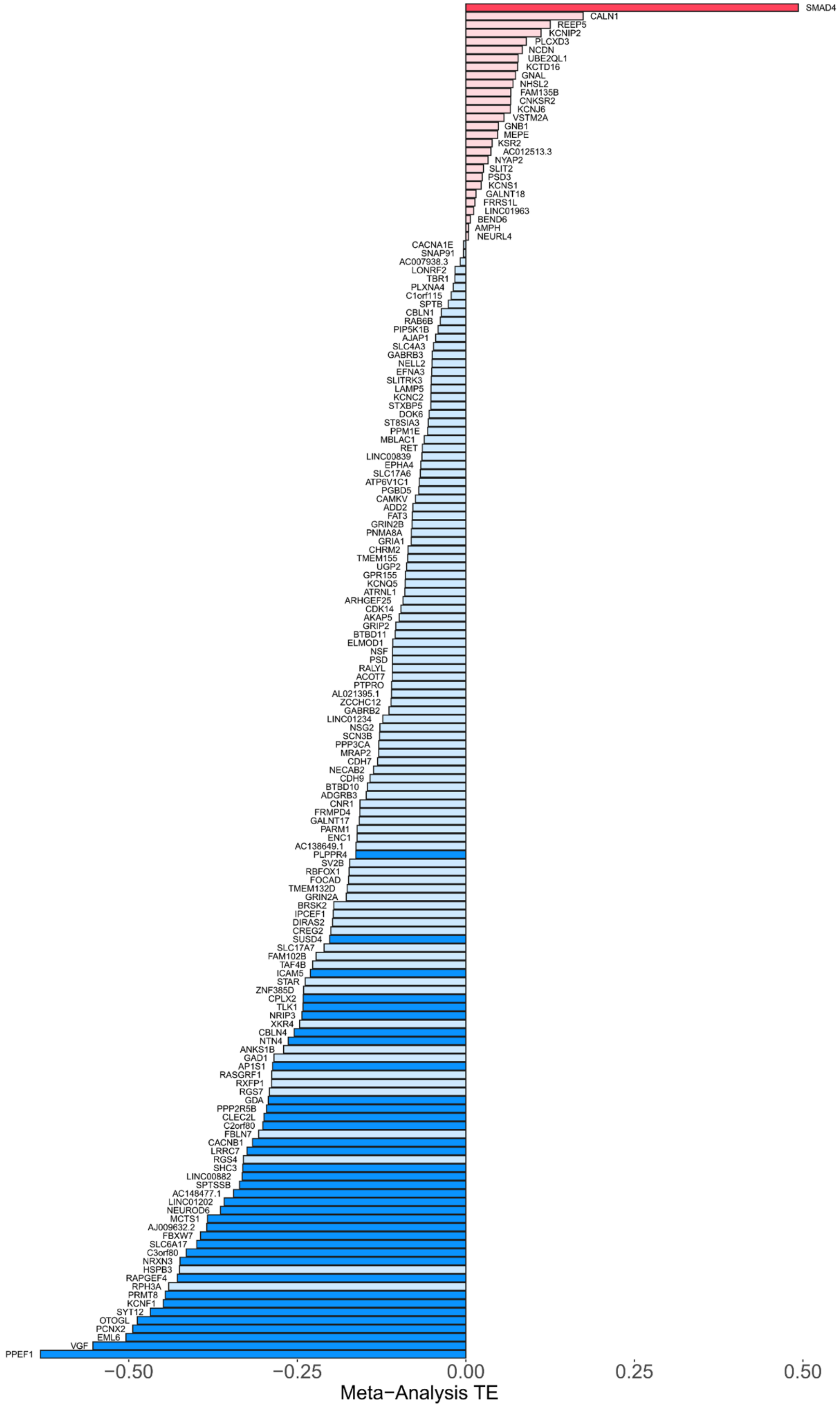


Waterfall plot showing PHGbrown subnetwork gene differential expression in AD cases versus controls, based on the published AMP-AD transcriptome meta-analysis. The color denotes down- (blue) or up- (red) regulated expression. Significant ($p_{meta}<0.05$) differentially expressed genes are indicated using darker shading.

**Fig. 9: Additional PHGbrown modifiers of adult brain degeneration.**

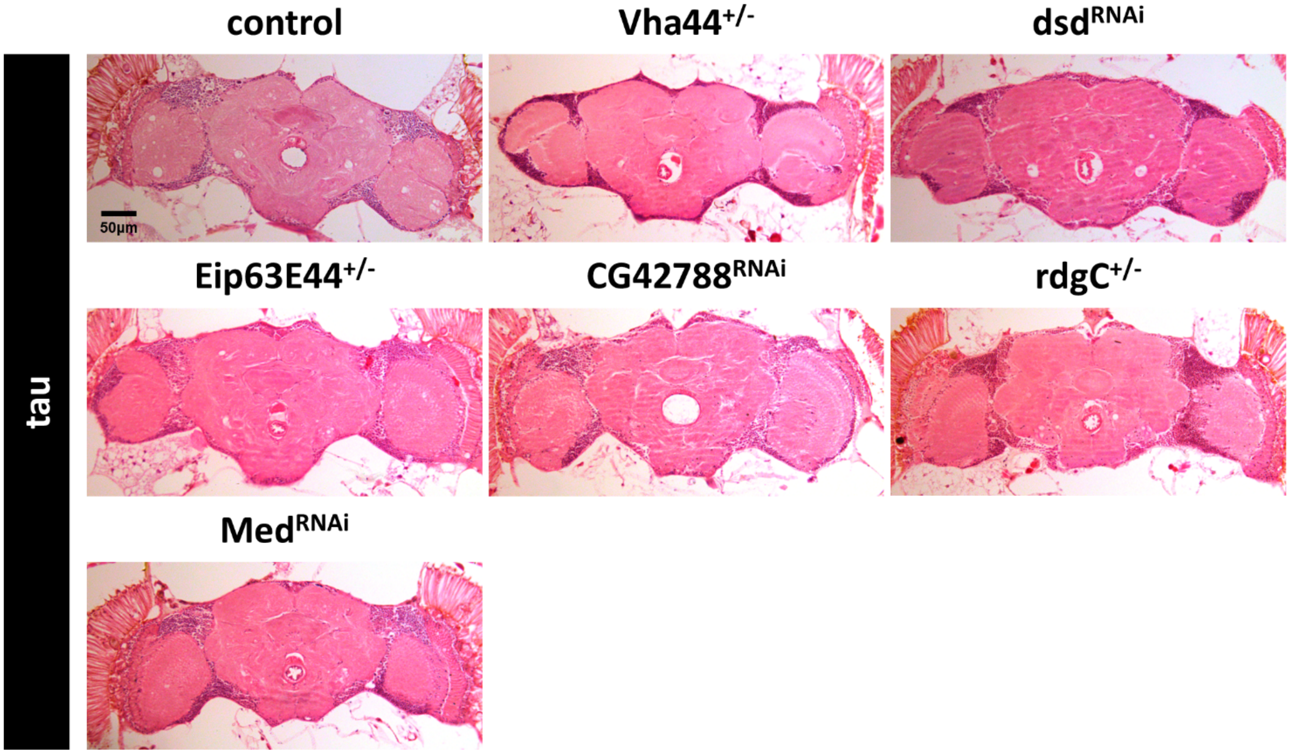


Additional homologs of PHGbrown causal drivers consistently modify tau induced neurodegeneration (*elav-Gal4/+; UAS-tau/+*) using an independent assay for vacuolar histopathology in the adult brain. The following RNAi knockdown or loss-of-function alleles were tested in heterozygosity: *Vha44(MI02871); dsd(v1106); Eip63E(MI00413-GFSTF.1); CG42788(v45034); rdgC(v35105); and Med(v19688).* See Fig. 5 for quantification and statistical analysis.

**Fig. 10: Additional studies of *VGlut* interactions with neurodegeneration.**

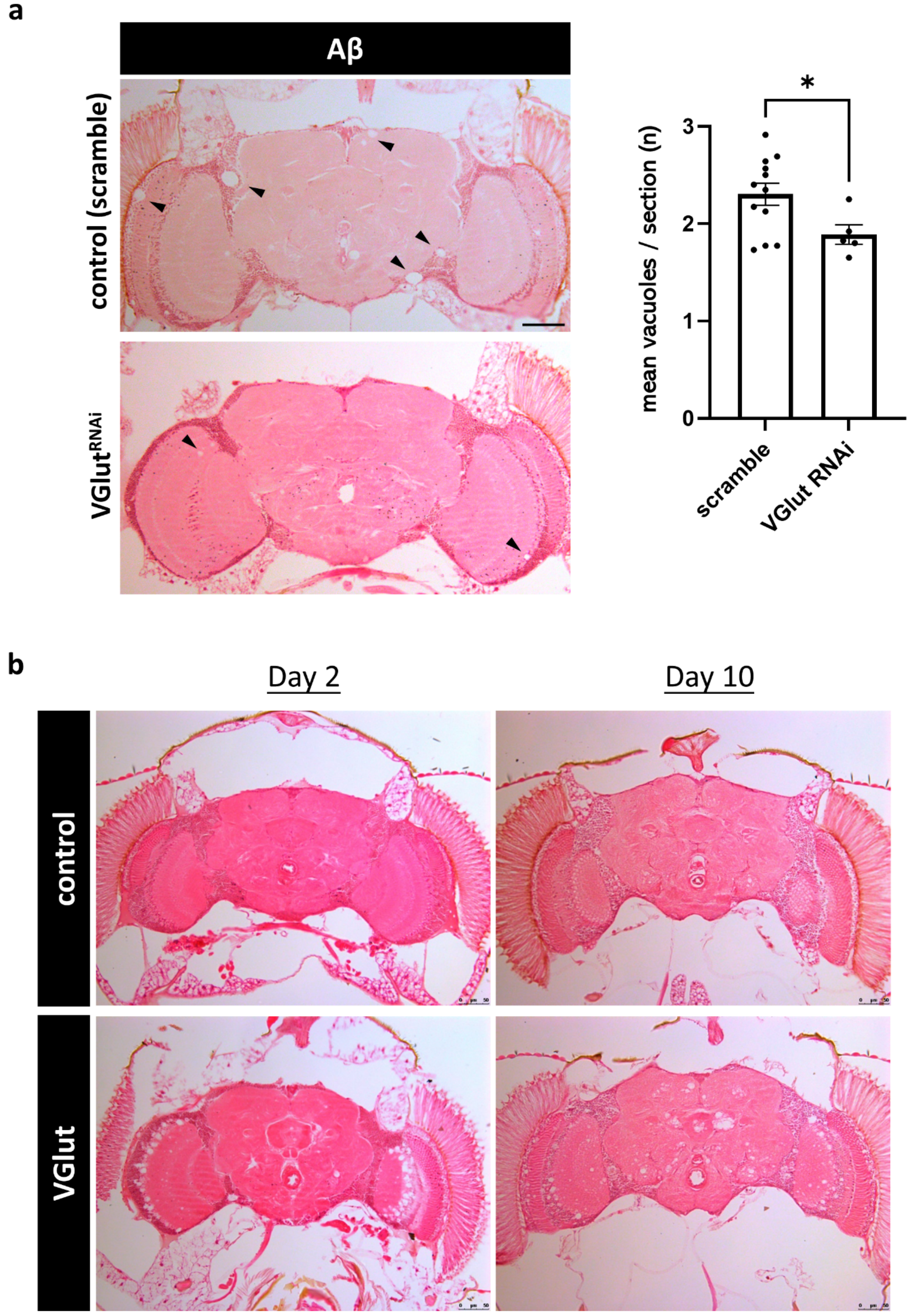


**a**, *VGlut* knockdown with RNA-interference (v2574) dominantly suppresses vacuolar degeneration caused by Aβ (*elav-Gal4/+; UAS-Aβ42/+*) in the adult *Drosophila* brain. The control expresses a "scramble" non-targeting RNAi transgene (v2691). Vacuoles (arrowheads) were quantified in at least n = 5 animals at 15 days of age. Statistical analysis based on two sample t-test. Each data point represents an individual biological replicate sample. Error bars denote the standard error of the mean. *, $p < 0.05$. Consistent results were obtained using a classical *VGlut* hypomorphic allele (see Fig. 6). **b**, *VGlut* overexpression (*VGlut-GAL4 / +; UAS-VGlut / +*) induces age-dependent, progressive vacuolar degenerative pathology, compared with driver controls (*VGlut-Gal4/+*).

**Fig. 11: Transcriptome analysis of the *VGlut* model of excitotoxic neurodegeneration.**

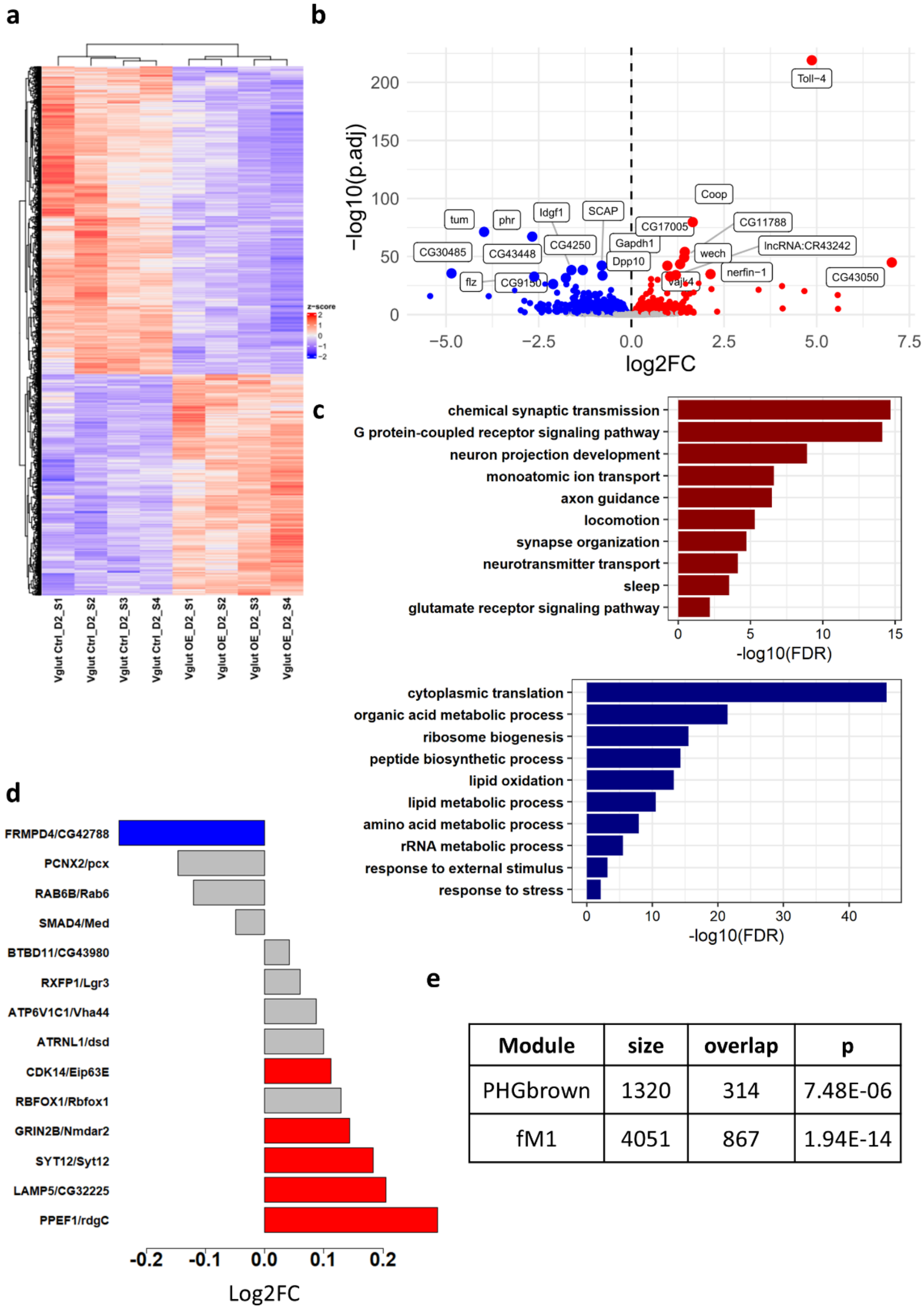


| Module | size | overlap | p |
|---|---|---|---|
| PHGbrown | 1320 | 314 | 7.48E-06 |
| fM1 | 4051 | 867 | 1.94E-14 |

**a**, RNA-sequencing was performed to evaluate brain gene expression changes in the *Drosophila VGlut* overexpression model. Heatmap showing z-scores of differentially expressed genes (rows) in each biological replicate (columns) including the *VGlut* overexpression model (*VGlut-Gal4 / +; UAS-VGlut / +*) and driver controls (*VGlut-Gal4 / +*). A total of n=4 replicates were analyzed for each experimental genotype in 2-day-old animals. Clustering used the default parameters of the pheatmap() function in R. **b**, Volcano plot showing the VGlut overexpression model RNA-seq differential expression results, displaying the magnitude of change ($log_2$-fold-change) versus statistical significance (negative log10 p value). Differentially expressed genes were defined based on a false discovery rate (FDR) adjusted significance level of $p < 0.05$. See also Extended Data Table 15. **c**, Gene ontology (GO) enrichment results of up- (red) and down- (blue) regulated diffrerentially expressed genes in comparisons of the *VGlut* overexpression model with driver controls. Selected terms reaching the significance level (FDR < 0.05) are shown. See also Extended Data Table 16. **d**, Many *Drosophila* homologs of PHGbrown driver gene modifiers are also dysregulated in the *VGlut* overexpression model of hyperactivation brain injury, with predominantly up-regulated expression. Red and blue bars denote significant differential expression (FDR adjusted p-value < 0.05). **e**, The transcriptional signature of *VGlut* overexpression (n=2,864 differentially expressed genes) significantly overlaps with the *Drosophila* fM1 synaptic module and PHGbrown (fly homologs), based on the hypergeometric overlap test.

**Fig. 12: Additional analysis of cell-type specific differential expression of PHGbrown.**

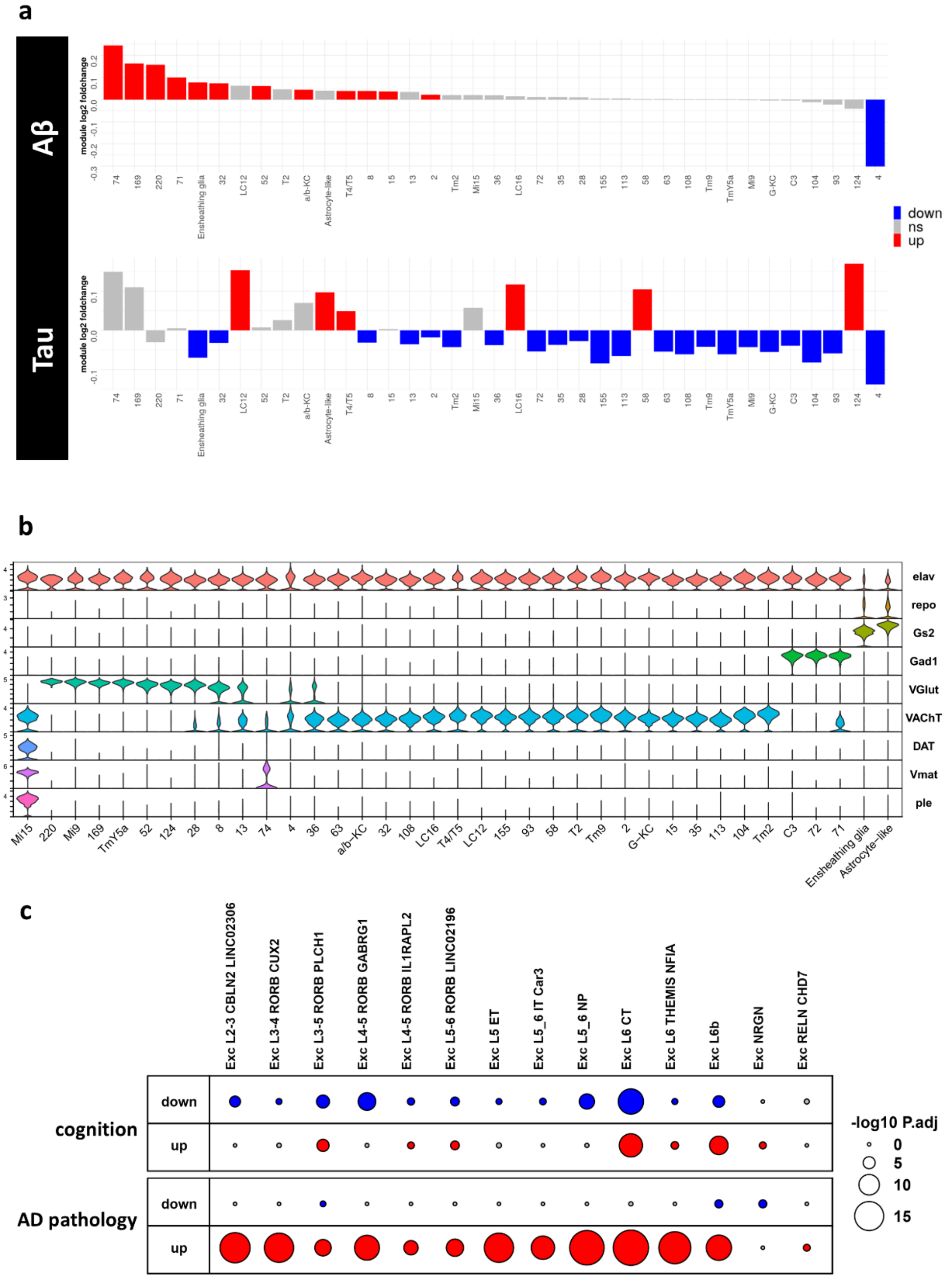


**a**, Cross-species analysis evaluating the expression of fly gene homologs of the human PHGbrown coexpression module in single-nucleus RNA-sequencing data from the Aβ (*elav-Gal4/+; UAS-Aβ42/+*) or tau (*elav-Gal4/+; UAS-tau/+*) models, compared with driver controls (*elav-GAL4/+*). Colored bars denote significant up (red) or down (blue) differential expression (Wilcoxon rank sum test $p<0.05$). See also Fig. 6 for complementary analysis of *Drosophila* fM1 module. **b**, Violin plot showing cell-type-specific marker gene expression across cell clusters evaluated from the *Drosophila* single nucleus RNA-seq data including *Elav* (neurons), *repo/Gs2* (glia), *Gad1* (GABA), *VGlut* (glutatamate), *VAChT* (acetylcholine), and *DAT/Vmat/ple* (dopamine). **c**, In human postmortem brain, PHGbrown is predominantly up-regulated in relation to AD pathology and cognitive decline in excitatory neuron populations. Differential expression analysis leveraged published AD single-nucleus RNA-sequencing data atlas from human postmortem brain tissue (n=427 brain autopsies) and considering global cognitive function at evaluations proximate to death (cogn_global_lv) and global AD pathology (gpath). Size of the circles correspond to significance of PHGbrown overlap based on the hypergeometric overlap test [-$\log_{10}$ (p-value), adjusted for false discovery rate], with signficant up and down regulated genes indicated in red versus blue, respectively. Of note, the clinical and pathologic traits selected (cogn_global_lv and gpath, respectively) are highly correlated but inversely related, with increasing pathologic burden associated with lower cognitive performance in evaluations proximate to death. See also Extended Data Table 17.
